# Contrasting population genetics of cattle- and buffalo-derived *Theileria annulata* causing tropical theileriosis

**DOI:** 10.1101/2020.01.10.902031

**Authors:** Umer Chaudhry, Qasim Ali, Lynn Zheng, Imran Rashid, Muhammad Zubair Shabbir, Muhammad Nauman, Kamran Ashraf, Mike Evans, Shahzad Rafiq, Muhammad Oneeb, Ivan Morrison, Neil D. Sargison

## Abstract

The present study was designed to improve understanding of *Theileria annulata* in sympatric water buffalo and cattle in the Punjab province of Pakistan. The prevalence of tropical theileriosis is high, buparvaquone resistance is widespread, and vaccine protection is poor in the field. Better understanding is, therefore, needed of the factors that influence the genetics of *T. annulata* populations both within its hosts and in its overall populations. Here we utilise a panel of six satellites and a mitochondrial cytochrome b marker to explore the multiplicity of *T. annulata* infection and patterns of emergence and spread of different parasite genotypes. Parasite materials were collected from infected animals in defined regions, where water buffalo and cattle are kept together. Our results show that *T. annulata* is genetically more diverse in cattle- than in water buffalo-derived populations (the mean numbers of unique satellite alleles were 13.3 and 1.8 and numbers of unique cytochrome b locus alleles were 65 and 27 in cattle- and water buffalo-derived populations, respectively). The data show a high level of genetic diversity among the individual host-derived populations (the overall heterozygosity (H_e_) indices were 0.912 and 0.931 in cattle, and 0.874 and 0.861 in buffalo, based on satellite and cytochrome b loci, respectively). When considered in the context of high parasite transmission rates and frequent animal movements between different regions, the predominance of multiple *T. annulata* genotypes, with multiple introductions of infection in the hosts from which the parasite populations were derived, may have practical implications for the spread of parasite genetic adaptations; such as those conferring vaccine cross-protection against different strains affecting cattle and buffalo, or resistance to antiprotozoal drugs.

## 1. Introduction

Vector-borne haemoprotozoa impact globally on the health, welfare, and production of livestock (Jabbar et al., 2015). The genus *Theileria* includes two particularly important species, *Theileria annulata* and *Theileria parva* (Lawrence, 1979). *T. parva* typically causes East Coast Fever, Corridor Disease, or January Disease in southern and eastern Africa (Uilenberg et al., 1982), whereas, *T. annulata* causes tropical theileriosis in North Africa and South Asia (Nourollahi-Fard et al., 2015). Tropical theileriosis is amongst the most important neglected tropical parasitic diseases of livestock (Sivakumar et al., 2014). Cattle and buffalo become infected with *T. annulata* following the transmission of sporozoites by ixodid ticks of the genus *Hyalomma*. These stages invade lymphocytes and develop into schizonts, which activate the infected cell to proliferate. Synchronous division of parasite and host cell results in rapid clonal expansion of the infected cell population. Some of the parasites undergo differentiation to produce merozoites, which are released from the lymphocytes enter erythrocytes where they develop to piroplasms. Piroplasms ingested during tick feeding, migrate to the arthropod gut where gametogeny and production of zygotes occurs. These enter the haemolymph and are carried to the salivary gland, where further intracellular development gives rise to sporozoites. The life cycle is completed when the next instar of the tick feeds (Gharbi and Darghouth, 2015). Clinically, infected cattle and buffalo show high fever and lymphadenopathy, sometimes accompanied by haemolytic anaemia, respiratory and ocular lesions (Al-Hosary et al., 2010; Mahmmod et al., 2011).

Measures used to control tropical theileriosis include avoidance, of infestation by *Hyalomma* ticks, including the application of acaricides, and prophylactic use of the theilericidal drug. However, these measures are expensive and difficult to apply. The unsustainability of these disease control measures highlights the potential for effective *T. annulata* vaccines. Attenuated live vaccines against *T. annulata* have been used in many countries where tropical theileriosis is endemic (Brown, 1990; Darghouth et al., 1999; Tait and Hall, 1990). However, there are practical constraints to the widespread use of live *T. annulata* vaccines in control programmes: (i) a requirement for distribution in liquid nitrogen, which accounts for approximately 30% of the cost (Bouslikhane et al., 1998); (ii) the need to use the vaccine immediately after thawing; (iii) the need for costly quality control measures to ensure consistent efficacy and the absence of other pathogens; (iv) in some cases, problems with post-vaccination reactions, which have been recorded in 3% of animals immunised with Chinese, Moroccan, Iranian or Tunisian stocks (Darghouth, 2008). To overcome these problems, research on the development of a killed subunit vaccine against *T. annulata* has focused on antigens present on the surface of sporozoite and merozoite stages of the parasite.

Genetic and antigenic diversity is a major constraint to the development of subunit vaccines for some parasite species. For example, cross-protection between cattle- and buffalo-derived *T. parva* strains is incomplete (Young et al., 1973). Studies of *T. parva* in African buffalo have shown a high level of genetic diversity when compared to cattle-maintained *T. parva* (Oura et al., 2005). While *T. annulata* can infect both cattle and Asian buffalo (Nourollahi-Fard et al., 2015), genetic comparisons between strains infecting sympatric host populations have not been reported. Understanding the genetic and antigenic diversity of *T. annulata* in buffalo and cattle and the extent to which the parasites circulate between the two species is needed to inform the potential applicability of vaccination.

Therapeutic control of *T. annulata* is heavily dependent on the use of a single theilericidal drug, namely buparvaquone. However, the effective use of this compound in some regions is now threatened by the emergence of drug resistance (Mhadhbi et al., 2015). Buparvaquone resistance has been reported in several countries and now represents a serious challenge to efficient livestock production (Mhadhbi et al., 2010). Although mutations in the candidate cytochrome b locus linked to resistance phenotype have been demonstrated, a causal relationship has not been established (Chatanga et al., 2019; Mhadhbi et al., 2015; Sharifiyazdi et al., 2012). Previous studies have described multiple genotypes and high levels of genetic diversity in *T. annulata* (Al-Hamidhi et al., 2015; Weir et al., 2011), although there is a lack of information on the genotypic diversity of drug-resistant parasites and whether there are fitness costs associated with mutations of the cytochrome b locus. It is, therefore, important to understand the genetic diversity and population sub-structure of *T. annulata*, with reference to the emergence and spread of buparvaquone resistance mutations.

Population genetic studies of *T. annulata* have previously been performed using panels of satellite DNA markers (Weir et al., 2007). More recently, deep amplicon sequencing of meta-barcoded DNA has afforded a practical and high-throughput method for investigating the genetic diversity between and within parasite populations. The Illumina MiSeq platform can provide 100,000 or more reads of up to 600 bp of loci of interest, depending on primer design. Recently, we have used these methods to study the population genetics of *Calicophoron daubneyi* (Sargison et al., 2019) and *Fasciola gigantica* infection in livestock keep in the United Kingdom and Pakistan.

In this paper, we describe the results of an analysis of the genetic diversity of buffalo- and cattle-derived *T. annulata,* by applying typing with six polymorphic satellite markers, combined with the deep amplicon sequencing of the mitochondrial cytochrome b locus. The results provide a comparison of the molecular profile of parasites in the two host species, which indicate greater genetic diversity in cattle parasites. These data represent an essential basis for further analyses to examine the emergence and spread of buparvaquone resistance in *T. annulata*.

## 2. Materials and Methods

### 2.1. Parasite resources, gDNA isolation and species identification

Four infected cell lines produced by the infection of blood mononuclear cells with laboratory-maintained stocks of *T. annulata* held in the University of Edinburgh, originally isolated in Turkey, India, Tunisia, and Morocco (Katzer et al., 1994), were used as reference controls. Blood samples were collected from the piroplasm-positive cattle and buffalo in veterinary clinics throughout the Punjab province of Pakistan between 2017 and 2019. Venous blood samples were collected into EDTA tubes were stored at −20°C. Samples were collected by para-veterinary staff under the supervision of local veterinarians following consent from the animal owners. The study was approved by the Institutional Review Board of the University of Veterinary and Animal Sciences (UVAS-24817). Peripheral blood smears were prepared and stained with 4% Giemsa, and examined microscopically to detect piroplasms. Genomic DNA was isolated from positive samples by lysis with GS buffer containing proteinase K as described in the TIANamp Blood DNA Kit (TIANGEN Biotech Co. Ltd, Beijing) and stored at −20°C. ‘Haemoprotobiome’ high-throughput sequencing (Chaudhry et al., 2019) was performed on piroplasm-positive blood samples to confirm the presence of *T. annulata*.

### 2.2. Adapter and barcoded PCR amplification of T. annulata cytochrome b

A 517 bp region of the cytochrome b locus was selected for deep amplicon sequencing. One μl gDNA of each of 31 buffalo, 54 cattle and the 4 positive control derived *T. annulata* samples was used as templates for the 1^st^ round adapter PCR amplification. A *de novo* primer set (Supplementary Table S1) was used under the following conditions: 13.25 μl ddH_2_O, 1 μl gDNA, 0.75 μl of 10 mM dNTPs mix, 5 μl of 5X HiFi Fidelity Buffer, 0.5 μl of 0.5U DNA polymerase enzyme (KAPA Biosystems, USA), and 0.75 μl of 10 μM forward adaptor primers and reverse adaptor primers. The thermocycling conditions were: 95°C for 2 min, 35 cycles at 98°C for 20 sec, 55°C for 15 sec, and 72°C for 2 min, followed by a final extension of 72°C for 2 min. The PCR product was purified with AMPure XP Magnetic Beads (1X) (Beckman coulter Inc., USA). A barcoded primer set was used in the 2^nd^ round of PCR amplification to add a fragment of unique sequence into each purified product (Supplementary Table S2) under the following conditions: 13.25 μl ddH_2_O, 2 μl bead purified PCR products, 0.75 μl of 10 mM dNTPs mix, 5 μl of 5X HiFi Fidelity Buffer, 0.75 μl of 0.5U DNA polymerase enzyme (KAPA Biosystems, USA), and 1.25 μl of 10μM forward and reverse primers. The thermocycling conditions were: 98°C for 2 min, 7 cycles at 98°C for 20 sec, 63°C for 20 sec, and 72°C for 2 min. The PCR products were purified with AMPure XP Magnetic Beads (1X) (Beckman coulter Inc., USA).

### 2.3. Illumina Mi-Seq run and data handling

A pooled library was prepared with 10 μl of barcoded bead purification PCR product from each *T. annulata* sample and sent to Edinburgh Genomics, UK for deep amplicon sequencing. The size of the amplicon was measured by qPCR library quantification (KAPA Biosystems, USA), before running on an Illumina Mi-Seq sequencer using a 600-cycle pair-end reagent kit (Mi-Seq Reagent Kits v2, MS-103-2003) at a concentration of 15 nM with addition of 15% Phix control v3 (Illumina, FC-11-2003). The Illumina Mi-Seq post-run processing uses the barcoded indices to split all sequences by sample and generate FASTQ files. These were analysed using Mothur v1.39.5 software (Schloss et al., 2009) with modifications in the standard operating procedures of Illumina Mi-Seq (Kozich et al., 2013) in the Command Prompt pipeline. Briefly, the raw paired read-ends were run into the ‘make.contigs’ command to combine the two sets of reads for each sample. The command extracted sequence and quality score data from the FASTQ files, creating the complement of the reverse and forward reads, and then joining the reads into contigs. After removing the too long, or ambiguous sequence reads, the data were aligned with the *T. annulata* cytochrome b reference sequence library prepared from the positive controls (for more details Supplementary Data S1 and section 2.1) using the ‘align.seqs’ command. Any sequences that did not match with the *T. annulata* cytochrome b reference library were removed and the ‘summary.seqs’ command was used to summarise the 517 bp sequence reads of the *T. annulata* cytochrome b locus. The sequence reads were further run on the ‘screen.seqs’ command to generate the *T. annulata* cytochrome b FASTQ file. Once the sequence reads were classified as *T. annulata*, a count list of the consensus sequences of each population was created using the ‘unique.seqs’ command. The count list was further used to create FASTQ files of the consensus sequences of each population using the ‘split.groups’ command (for more details Supplementary Data S2).

### 2.4. Bioinformatics data analysis

The consensus sequences of *T. annulata* cytochrome b locus were aligned using the MUSCLE alignment tool in Geneious v10.2.5 software (Biomatters Ltd, New Zealand) and then imported into the FaBox 1.5 online tool (birc.au.dk) to collapse the sequences that showed 100% base pair similarity after corrections into a single genotype. The genotype frequency of each sample was calculated by dividing the number of sequence reads by the total number of reads. A split tree was created in the SplitTrees4 software (bio-soft.net) by using the neighbour-joining method in the JukesCantor model of substitution. The appropriate model of nucleotide substitutions for neighbour-joining analysis was selected by using the jModeltest 12.2.0 program (Posada, 2008). The tree was rooted with the corresponding cytochrome b sequence of *T. parva*. The branch supports were obtained by 1000 bootstraps of the data. The genetic diversity of cytochrome b was calculated within and between populations by using the DnaSP 5.10 software package (Librado and Rozas, 2009), and the following values were obtained: Heterozygosity (H_e_), the number of segregating sites (S), nucleotide diversity (π), the mean number of pairwise differences (k), the mutation parameter based on an infinite site equilibrium model, and the mutations parameter based on segregating sites (S_θ_).

### 2.5. Satellite genotyping and bioinformatics data analysis of T. annulata

Six satellite markers (TS6, TS8, TS20, TS31, TS12, TS16) were selected for use on each population as previously described by Weir et al. (2007). A summary of primer sequences and allele ranges is given in Supplementary Table S3. Eighteen buffalo and 35 cattle derived *T. annulata* populations were successfully analysed from six markers. PCR amplification was performed using a master mix containing 17.2 μl ddH_2_O, 0.3 μl of 100 μM dNTPs, 2.5 μl of 1X thermopol reaction buffer, 0.3 μl of 1.25U Taq DNA polymerase (New England Biolabs) and 0.3 μl of 0.1μM forward and reverse primers. The thermo-cycling parameters were 94°C for 2 min followed by 35 cycles of 94°C for 1 min, staying at the annealing temperatures of 60°C (TS6, TS20), 55°C (TS8, TS12, TS16), or 50°C (TS31) for 1 min and 65°C for 1 min, with a single final extension cycle of 65°C for 5 min. The forward primer of each microsatellite primer pair was 5’ end labelled with fluorescent dye (IDT, UK), and the ROX 400 internal size standard was used on the ABI Prism 3100 genetic analyser (Applied Biosystems, USA).

Individual chromatograms were analysed using Peak Scanner software version 2.0 (Thermo Fisher Scientific, USA) to determine the size of the alleles. These were combined across six markers to generate a multilocus genotype (MLG). From the MLG data, allelic variation (A) and heterozygosity (H_e_) for individual populations were calculated using Arlequin 3.11 software (Guo and Thompson, 1992). Significance levels were calculated using the sequential method of Bonferroni correction for multiple comparisons in the same dataset (Rice, 1989). Analysis of molecular variance (AMOVA) was estimated through the partition of satellite diversity between and within populations (Excoffier et al., 1992) and fixation index (pairwise F_ST_) values were calculated using Arlequin 3.11 to provide a measurement of population genetic sub-structure. Principal coordinate analysis (PCoA) was performed using GenALEx software to illustrate the extent of genetically distinct features of individual populations with plot coordinates (Peakall and Smouse, 2012).

The likelihood ratio test statistics (G-test) were calculated using Arlequin 3.11 software (Excoffier et al., 2005) to estimate genetic linkage equilibrium and deviations from Hardy-Weinberg equilibrium. There was no evidence to support linkage disequilibrium for any combination of satellite loci in the individual *T. annulata* populations of buffalo and cattle, indicating that alleles at these loci were randomly associating and not genetically linked (data on file). There was some significant departure from Hardy-Weinberg equilibrium, even after Bonferroni correction, in addition to relative P values for 108 loci combinations for buffalo and 210 loci combinations for cattle *T. annulata* populations. The bottleneck software (version 1.2.02) was, therefore, used to search for the evidence of heterozygosity excess and mode-shift. The Wilcoson signed-rank test was used to evaluate the statistical significance of any possible genetic drift equilibrium (Lefterova et al., 2015). The data showed that there was no heterozygosity excess according to the Sign Test and Wilcoxon Test (data on file). The mode shift analysis established most populations had a normal L-shaped distribution.

## 3. Results

### 3.1. Allelic diversity between cattle- and buffalo-derived T. annulata populations

The allelic diversity data show that *T. annulata* is more diverse in cattle as compared to water buffalo. The mean numbers (±SD) of satellite alleles in 35 cattle- and 18 buffalo-derived populations were 34.7 (±9.1) and 23.7 (±5.2), respectively. The number of alleles (A_n_) for each satellite marker in the cattle- and buffalo-derived populations ranged from 22 to 46 and 17 to 32, respectively. The mean numbers (±SD) of cattle- and buffalo-specific *T. annulata* satellite alleles (A_u_) per marker were 13.3 (±6.3) and 1.8 (±2.04), respectively (Table 1 and Fig. 1). Seventy-nine and 41 cytochrome b locus alleles (A_n_) were identified in the 54 cattle- and 31 buffalo-derived *T. annulata* populations. The numbers of cattle- and buffalo-specific *T. annulata* alleles (A_u_) were 65 and 27, respectively (Table 1 and Fig. 1). The high level of allelic variation described in *T. annulata* might have practical implications for the development of a sporozoite and merozoite antigen subunit vaccine and immunisation cross-protection.

**Fig. 1.**
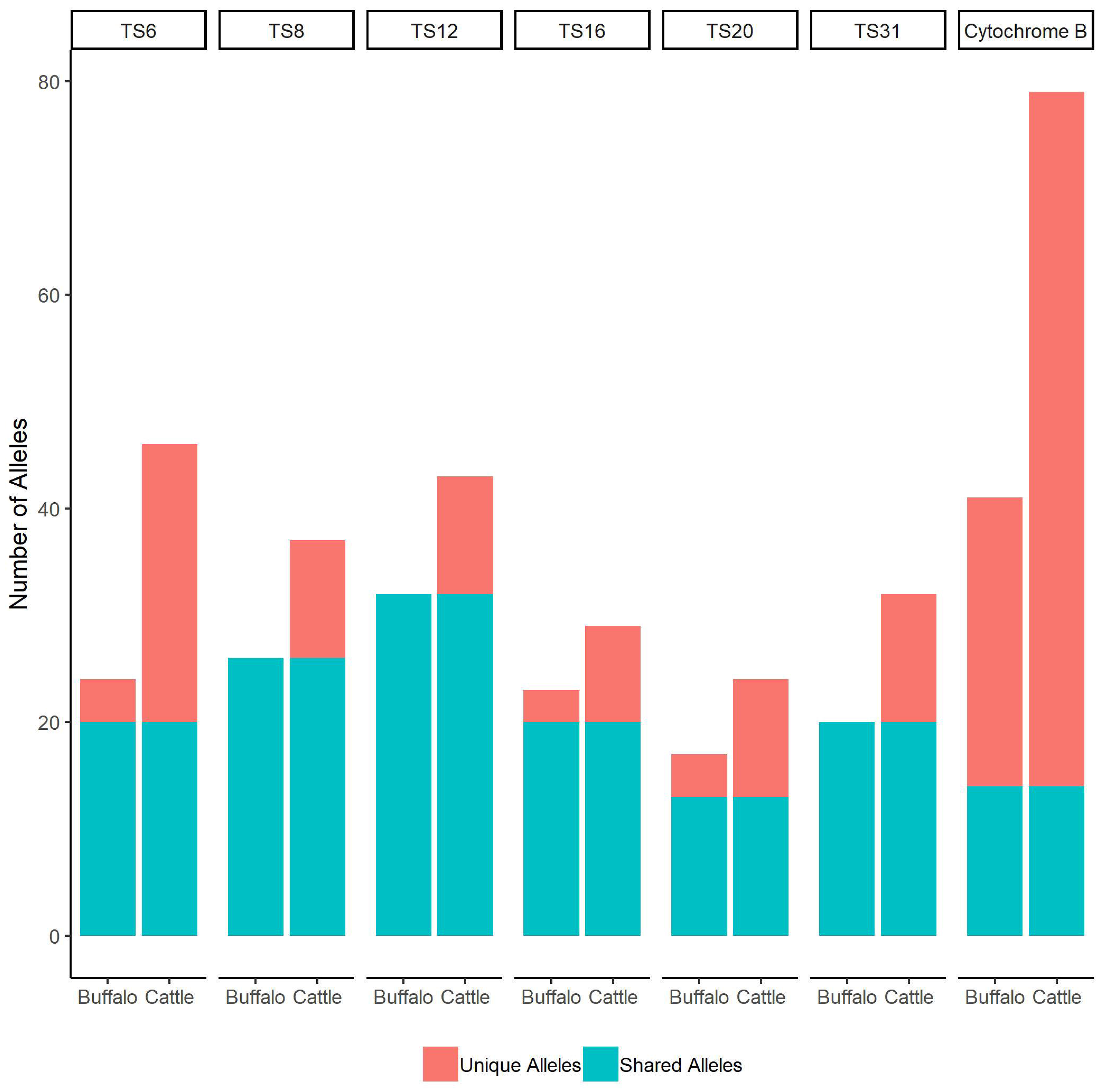
The mean number of alleles in *T. annulata* derived from buffalo and cattle based on a panel of six satellite and a cytochrome b marker. The bar of each marker shows the proportion of alleles in buffalo and cattle. The Y-axis shows the number of alleles of each marker. The unique and shared alleles of buffalo and cattle are represented by a different colour.

### 3.2. Genetic diversity within T. annulata populations derived from individual cattle and buffalo

The heterozygosity (H_e_) data reveal high levels of genetic diversity within each individual populations (35 cattle and 18 buffalo) of *T. annulata*; with mean values (±SD) for the six satellite markers ranging from 0.8 (±0.082) to 0.959 (±0.017) in cattle- and 0.738 (±0.196) to 0.956 (±0.034) in buffalo-derived populations (Table 2). Overall, the mean heterozygosities (H_e_) (±SD) of the cattle- and buffalo-derived *T. annulata* populations were 0.912 (±0.056) and 0.874 (±0.028) respectively (Table 2). High levels of heterozygosity (H_e_) were seen at the *T. annulata* cytochrome b locus, ranging from 0.556 to 0.923 (overall mean value 0.931) in each of the 54 cattle-derived populations and from 0.545 to 0.900 (overall mean value 0.861) in each of the 31 buffalo-derived populations (Table 3). The high level of genetic diversity in *T. annulata* may have practical implications for the emergence of buparvaquone drug resistance mutations.

### 3.3. Genetic sub-structure and phylogenetic relationship between T. annulata populations derived from individual cattle and buffalo

Genetic differentiation was shown by comparing the fixation indices (F_ST_) of two populations with each other in a pairwise manner. The F_ST_ values indicated a low level of genetic differentiation between *T. annulata* ranging from 0.001 to 0.114 in each of the 35 cattle-derived populations and 0.003 to 0.132 in each of the 18 buffalo-derived populations, respectively, respectively (Supplementary Table S4). An AMOVA was conducted using the panel of six satellite markers to estimate the genetic variation within and between the populations. This showed that genetic variation was distributed 97.91% and 97% and within; and 2.09% and 3% between cattle- and buffalo-derived *T. annulata* populations, respectively (data on file). PCoA was performed to illustrate as a two-dimensional plot the extent to which populations are genetically distinct. The two axes accounted for 72.5% (25.67 + 46.83) and 73.93% (27.6 + 46.33) of the variation in the cattle- and buffalo-derived populations, respectively; and showed that populations from different regions formed overlapping clusters, hence were not geographically sub-structured (Fig. 2).

**Fig. 2.**
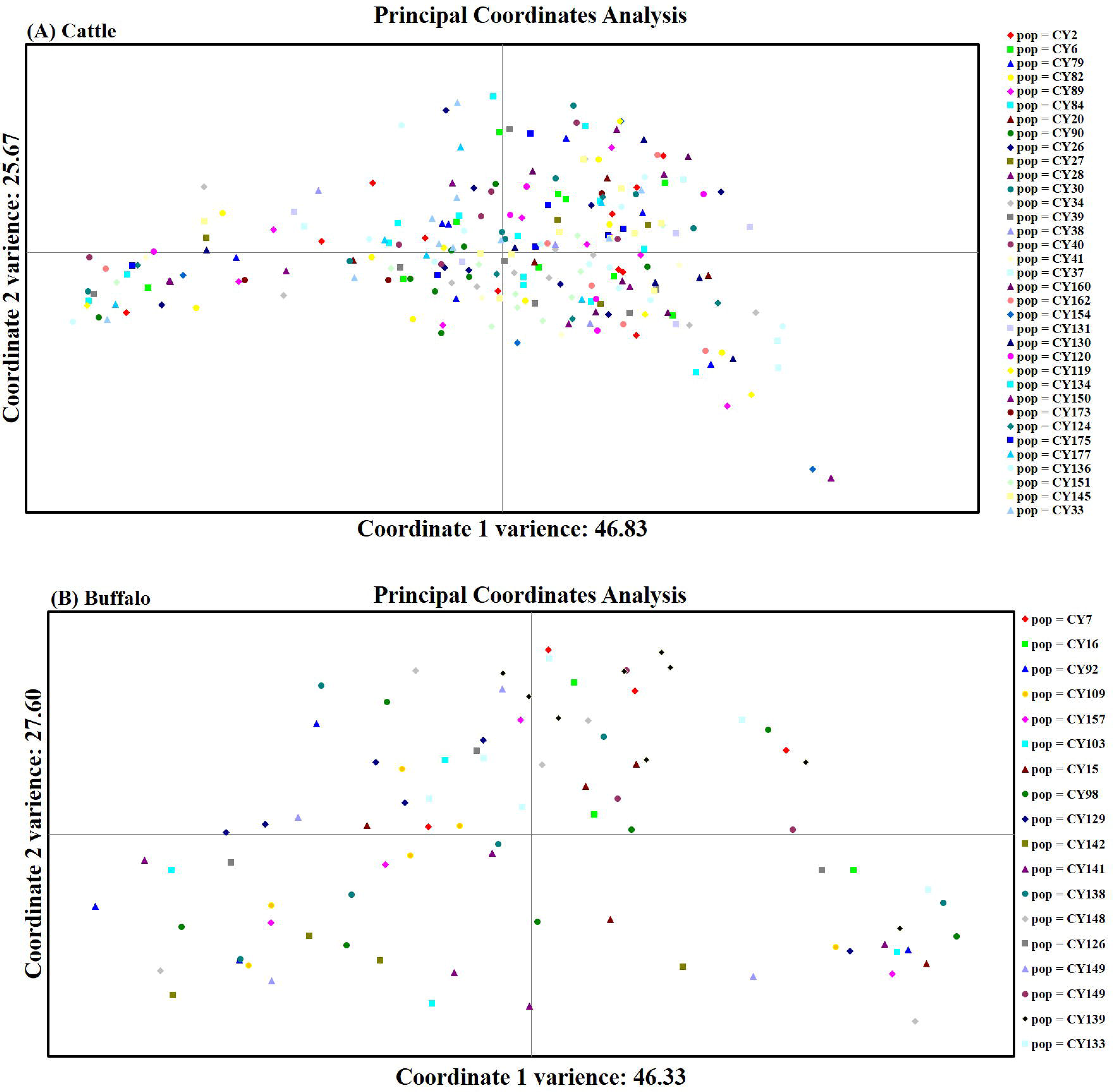
Principal coordinate analysis using a panel of six satellite markers to represent 35 cattle- and 18 buffalo-derived *T. annulata* populations.

Phylogenetic analysis of the seventy-nine cytochrome b genotypes of the 54 cattle-derived *T. annulata* populations, showed that two genotypes for 22.9% and 10.8% of the total number of the sequence reads, being present in 36 and 8, populations, respectively. Seven genotypes accounted for between 5.0% and 7.0% of the sequence reads and 45 genotypes accounted for less than 1.0% of the sequence reads (Fig. 3A). The split tree shows that eight genotypes are shared between populations derived from cattle holdings in multiple locations of Lahore, Chakwal, Gujranwala, Okara, and Sahiwal. Seven genotypes are shared between populations derived from cattle holdings in any two locations of Lahore, Gujranwala, Okara, and Sahiwal. Sixty-one genotypes are present in a single location (Gujranwala=24; Qadirabad=12; Okara =14; Lahore=11 genotypes) (Fig. 3A).

**Fig. 3.**
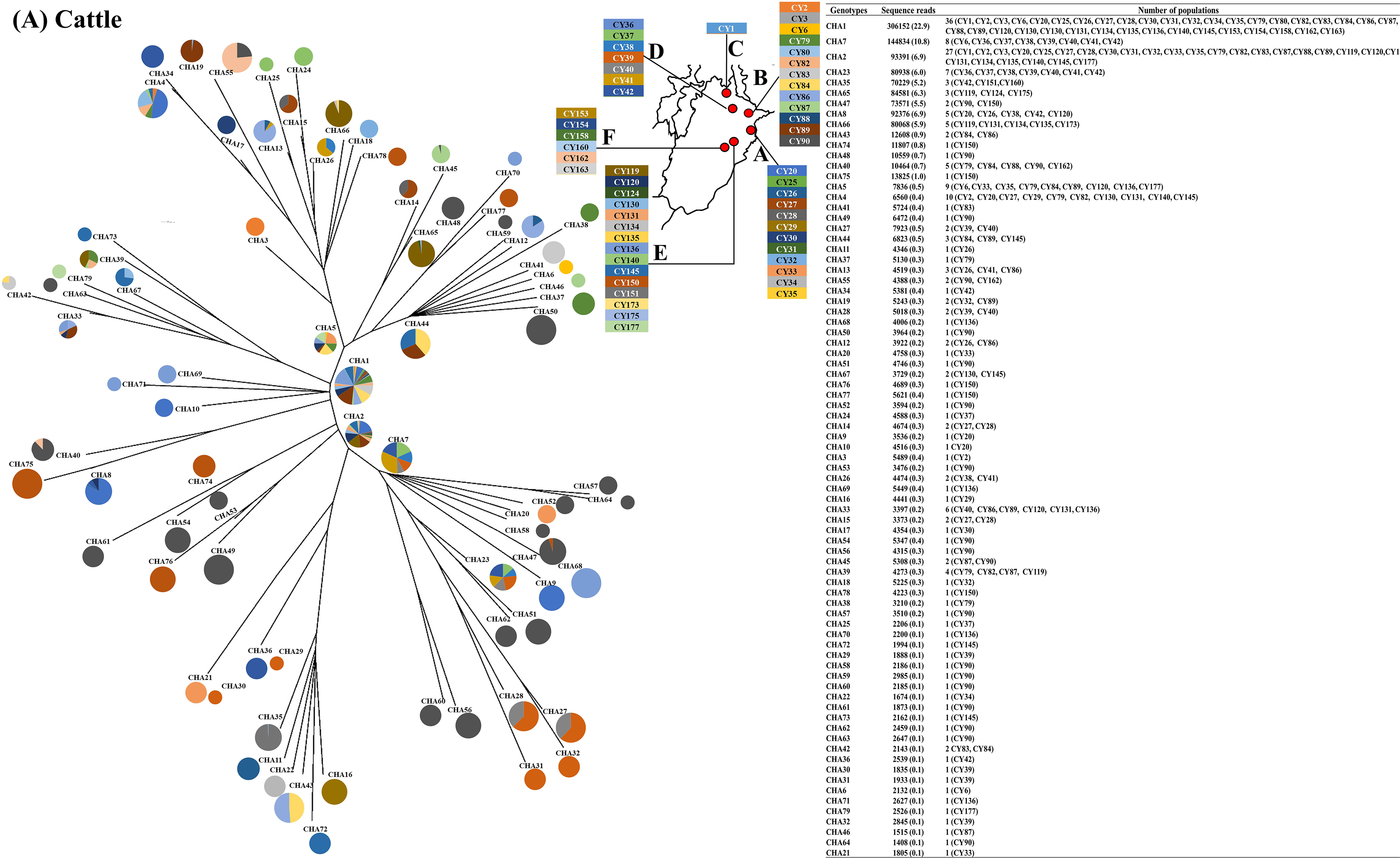
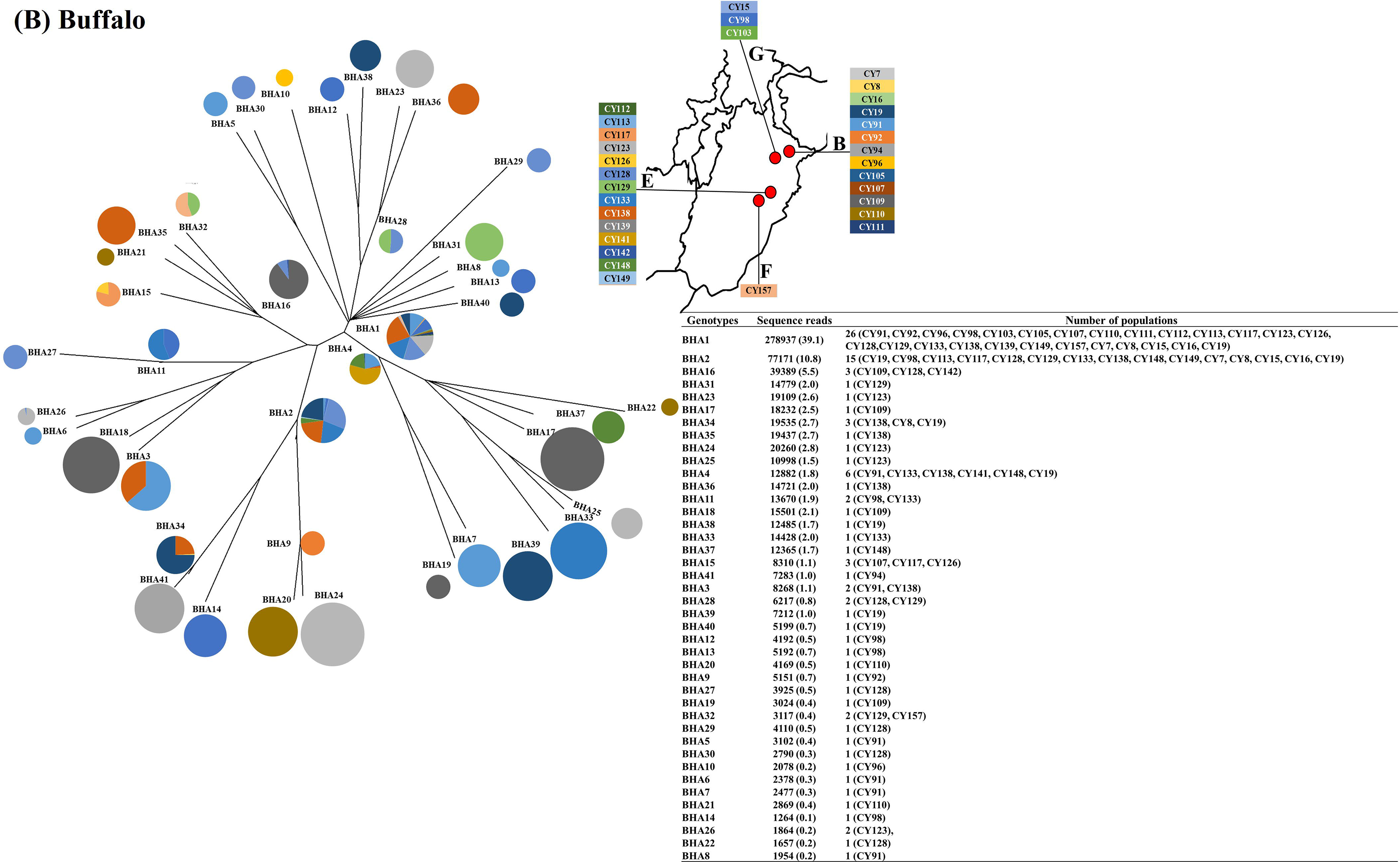
Split tree analysis of 79 cattle-derived (3A) and 41 buffalo-derived (3B) *T. annulata* genotypes collected from the Lahore (A), Gujranwala (B), Chakwal (C), Qadirabad (D), Okara (E), Sahiwal (F), and Hafizabad (G) regions of the Punjab province of Pakistan. Each population (CY) is represented by a different colour in the individual genotype (CHA and BHA). The pie chart circles represent the distribution and percentage of sequence reads generated per genotype identified in 54 cattle- and 31 buffalo-derived populations, as indicated in the insert table.

Phylogenetic analysis of the forty-one cytochrome b genotypes of the 31 buffalo-derived *T. annulata* populations, showed that three genotypes accounted for 39.1%, 10.8%, and 5.5% of sequence reads being present in 26, 15 and 3 populations, respectively. Eighteen genotypes accounted for between 1.0% and 3.0% of sequence reads and 20 genotypes accounted for less than 1.0% of sequence reads (Fig. 3B). The split tree shows that two genotypes are shared between *T. annulata* populations derived from buffalo holdings in multiple locations of Gujranwala, Hafizabad, Okara, and Sahiwal. Five genotypes are present in the populations derived from buffalo holdings in Gujranwala and Okara. One genotype is present in the populations derived from buffalo holdings in Hafizabad and Okara and one genotype from Okara and Sahiwal. Thirty-two genotypes are present in a single location (16 genotypes in Gujranwala; 13 genotypes in Okara; and 3 in Hafizabad) (Fig. 3B). The satellite and cytochrome b data provide evidence of a high level of gene flow predicted to occur due to livestock movements or translocations of ticks to a new region. This could potentially influence the spread of drug-resistant alleles.

### 3.4. Genotype distribution of T. annulata populations derived from individual cattle and buffalo

*T. annulata* satellite data were found to be highly polymorphic in each individual population (35 cattle and 18 buffalo), with overall numbers of genotypes per population ranging from 2 to 26 in cattle and 2 to 17 in buffalo. The mean numbers (±SD) of *T. annulata* genotypes in cattle- and buffalo-derived populations were 8.60 (±2.52) and 6.25, respectively (Table 4).

Seventy-nine *T. annulata* cytochrome b genotypes from 54 individual cattle-derived populations were analysed separately. Fifteen populations had a single genotype at high frequencies ranging from 90 to 100%. These comprised of ten populations containing only one genotype; two populations containing two genotypes; one population with three genotypes; one population with four genotypes; and one population with seven genotypes (Fig. 4A). Thirty-nine populations had multiple genotypes at high frequencies. These comprised of five populations containing two genotypes; eight populations containing three genotypes; ten population containing four genotypes; seven populations with five genotypes; four populations with six genotypes; one population with seven genotypes; two populations with eight genotypes; one populations with 12 genotypes; and one with 14 genotypes (Fig. 4A).

**Fig. 4.**
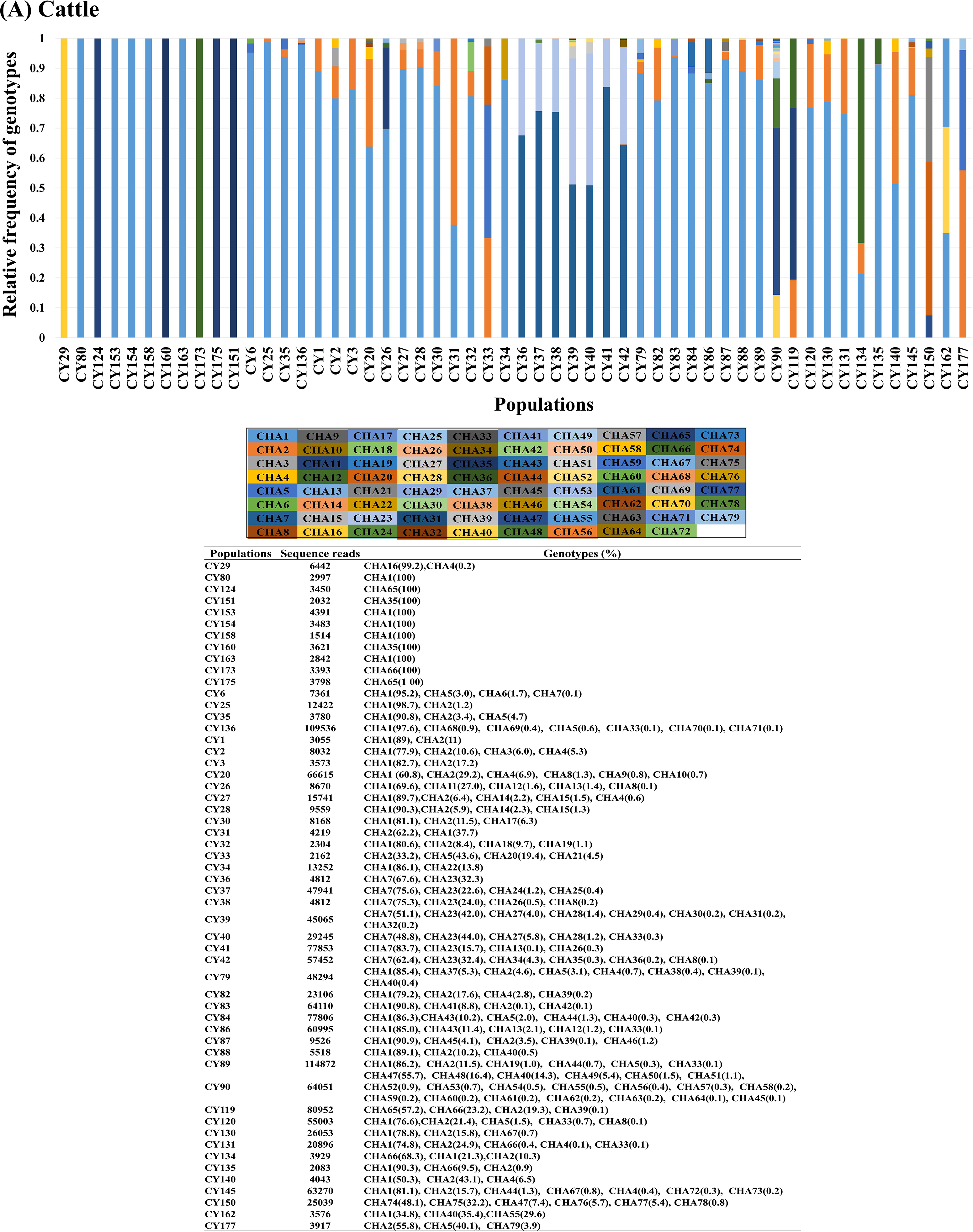
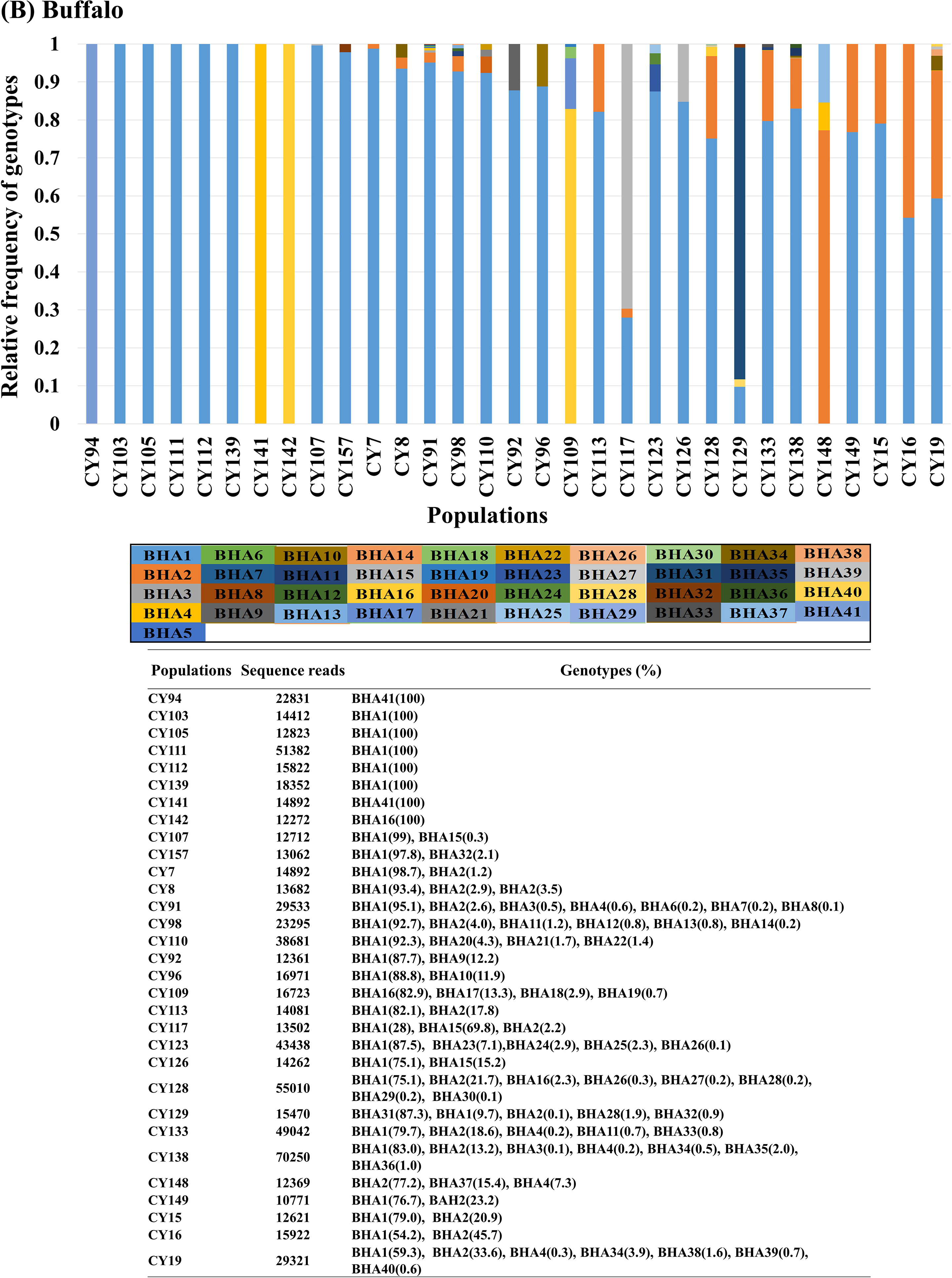
Relative genotype frequencies of the cytochrome b locus of 54 cattle-derived (4A) and 31 buffalo-derived (4B) *T. annulata* populations, collected from the Punjab province of Pakistan. Each genotype (CHA and BHA) is represented by a different colour in the individual population (CY). The population distribution and the frequency of sequence reads generated per population for each of the 79 cattle-derived and 41 buffalo-derived *T. annulata* genotypes is shown in the insert table.

Forty-one *T. annulata* cytochrome b genotypes from 31 individual buffalo-derived populations were analysed separately. A single genotype predominated in 15 populations at a frequency of between 90 and 100%. These comprised of eight populations containing only one genotype; three populations containing two genotypes; one population with three genotypes; one population with five genotypes; one population with six genotypes; and one population with seven genotypes (Fig. 4B). Sixteen populations had high frequencies of multiple genotypes. These comprised of seven populations contained 2 genotypes; two populations contained 3 genotypes; one population contained 4 genotypes; three populations had 5 genotypes; two populations had 7 genotypes and one population contained a maximum of 8 genotypes (Fig. 4B). The satellite and cytochrome b data provide evidence of the potential impact of the multiplicity of *T. annulata* infection and their control strategies for tropical theileriosis.

## 4. Discussion

*Theleria annulata* is considered to be the most economically important protozoan parasite of cattle and water buffalo in South Asia, the Middle East and North Africa, where livestock production is critically important. *T. annulata* has a major impact on food production in low- and middle-income countries (LMICs), where efficient agriculture is a priority and the disease causes economic losses due to high mortality and morbidity of livestock, having a significant impact on meat and milk production (Jabbar et al., 2015). Our results reveal a high level of genetic diversity within *T. annulata* infecting individual hosts, and between different host-derived populations. Although both cattle and buffalo parasite populations showed extensive diversity, the extent of diversity was significantly greater in cattle compared to buffalo. The diversity and distribution of *Theileria annulata* genotypes in the Punjab province of Pakistan are consistent with high parasite transmission rates and frequent animal movements within the region.

The satellite and cytochrome b data reported in the present study show that *T. annulata* is genetically more diverse, with more circulation of alleles, in cattle as compared to water buffalo. The satellite data were informative due to the numbers of alleles present, showing more unique alleles in cattle-derived than in Asian water buffalo-derived *T. annulata* populations. The cytochrome b locus, although genotypically less polymorphic, provided further evidence of allelic diversity and differences between cattle- and buffalo-populations. Similarly, high levels of parasite heterogeneity have been reported in *T. parva*, although the diversity in cattle is much more restricted compared to the buffalo host (Oura et al., 2011). In *T. parva*, there is evidence that although most if not all buffalo-derived parasites are capable of infecting cattle, many are incapable of undergoing tick transmission. Hence, differences in the capacity of different *T. annulata* genotypes to undergo tick transmission in cattle and buffalo could potentially influence the genotypic composition of the parasites that are maintained in the two host species. Immunisation of cattle with different subunit versions of the sporozoite stage surface protein (SPAG1) has provided a degree of protection against cattle-derived *T. annulata*, probably by neutralising the ability of the parasite to infect host cells (Schnittger et al., 2002). Given the high overall level of genetic diversity in *T. annulata*, it will be important to determine whether there is polymorphism in candidate antigens that would influence the ability to generate broadly cross-reactive immunity (Katzer et al., 1994).

Our satellite and cytochrome b locus data shows a high levels of genetic diversity in both cattle- and buffalo-derived *T. annulata* populations, reflecting high effective population sizes (Al-Hamidhi et al., 2015). *T. annulata* genome comprises 8.35 Mb spanning four chromosomes that range from 1.9 to 2.6 Mb, with 3,792 putative protein-coding genes. In addition, a total of 49 transfer RNA and five ribosomal RNA genes were identified (Pain et al. (2005). High level of genetic diversity in *T. annulata* rise the possibility of the emergence of potential drug resistance mutations. Pakistani livestock is treated frequently with buparvaquone and anecdotal evidence suggests that resistance is widespread, implying that there is likely to be strong selective pressure for the emergence of drug resistance. The importance of this point is emphasised by the fact that buparvaquone is the only drug available for the treatment of tropical theileriosis.

Fifteen cytochrome b genotypes were shared between the *T. annulata* populations of cattle and nine were shared between cattle populations. The satellite data provide further evidence of low levels of genetic differentiation among cattle-and buffalo-derived populations and show overlapping clusters that are not geographically sub-structured, consistent with the high levels of gene flow due to livestock movements or translocations of ticks to a new region. This could influence the spread of drug-resistant alleles. Studies of the global genetic sub-structure of *T. annulata* have shown a high level of genetic differentiation between Turkey and Tunisia (Weir et al., 2007). Studies have also shown a low level of genetic differentiation in *T. annulata* within-countries including Turkey, Tunisia, Oman, China and Portugal (Al-Hamidhi et al., 2015; Gomes et al., 2016; Weir et al., 2011; Yin et al., 2018), reflecting high levels of animal movement, or translocation of ticks.

The presence of a single cytochrome b genotype in 15 cattle and 9 buffalo- derived populations suggests a single emergence of *T. annulata* infection. In contrast, the predominance and the high proportions of multiple cytochrome b genotypes in 39 cattle- and 16 buffalo-derived populations implies multiple emergences of infection. The numbers of satellite genotypes per population provide further evidence for the pattern of the multiplicity of infection in genetically different adapted strains. Random cross mating of gametes and genetic recombination of *T. annulata* in ticks gives rise to the formation of new genotypes (Al-Hamidhi et al., 2015). The multiplicity of *T. annulata* infection may be influenced by variations in the intensity of transmission due to levels of different tick species infestation, or prevalence of tick infection (Yin et al., 2018). *Hyalomma scupense* is the most common and economically important species and the major vector of *T. annulata*, but *Hyalomma marginatum* and *Hyalomma anatolicum* are also found in Tunisia (Bouattour et al., 1996). There are four *Hyalomma* species responsible for the transmission of *T. annulata* in Turkey, of which *H. anatolicum* is the major vector for *T. annulata* (Aktas et al., 2004). Two tick species, *Hyalomma lusitanicum*, and *H. marginatum* are the vectors of *T. annulata* in Portugal (Estrada-Pena and Santos-Silva, 2005). There are a few reports that *H. anatolicum* may be a vector of *T. annulata* in the Punjab province of Pakistan (Karim et al., 2017). The multiplicity of *T. annulata* infection may also be influenced by population bottlenecking effects, arising from seasonal effects of climatic conditions on tick transmission and completion of the parasite’s lifecycle. The *T. annulata* transmission season in Turkey is between May and September, with the peak of clinical cases occurring in mid-summer (Sayin et al., 2003). In a region of endemic stability of tropical theileriosis in Tunisia, there is a high level of multiplicity of *T. annulata* infection, but clinical disease is rare; while in a region of endemic instability, a proportion of the population becomes infected all year round, and the clinical disease occurs particularly in adult cattle (Gharbi et al., 2011).

## 5. Conclusions

Control strategies for tropical theileriosis need to consider factors such as: the reproductive isolation of parasite populations; management of host movements; control of tick vectors; mitigation of the impacts of climate change; and consequences of parasite exposure to antiprotozoal drugs. Understanding of the multiplicity of infection and of high levels of gene flow is, therefore, important in the educational dissemination and implementation of advice on sustainable parasite control.

## Acknowledgement

The study was financially supported by the Carnegie Trust Scotland and Biotechnology and Biological Sciences Research Council (BBSRC). Work at the University of Veterinary and Animal Science Pakistan uses facilities funded by the Higher Education Commission of Pakistan. The authors of this study would like to thank Vice-Chancellor of the University of Veterinary and Animal Science Lahore Pakistan, for his great support in the arrangements for sample collections.

## Conflict of interest

None

## Notes

#### Summary of Updates

The present study was designed to improve understanding of Theileria annulata in sympatric water buffalo and cattle in the Punjab province of Pakistan. The prevalence of tropical theileriosis is high, buparvaquone resistance is widespread, and vaccine protection is poor in the field. Better understanding is, therefore, needed of the factors that influence the genetics of T. annulata populations both within its hosts and in its overall populations. Here we utilise a panel of six satellites and a mitochondrial cytochrome b marker to explore the multiplicity of T. annulata infection and patterns of emergence and spread of different parasite genotypes. Parasite materials were collected from infected animals in defined regions, where water buffalo and cattle are kept together. Our results show that T. annulata is genetically more diverse in cattle- than in water buffalo-derived populations (the mean numbers of unique satellite alleles were 13.3 and 1.8 and numbers of unique cytochrome b locus alleles were 65 and 27 in cattle- and water buffalo- derived populations, respectively). The data show a high level of genetic diversity among the individual host-derived populations (the overall heterozygosity (He) indices were 0.912 and 0.931 in cattle, and 0.874 and 0.861 in buffalo, based on satellite and cytochrome b loci, respectively). When considered in the context of high parasite transmission rates and frequent animal movements between different regions, the predominance of multiple T. annulata genotypes, with multiple introductions of infection in the hosts from which the parasite populations were derived, may have practical implications for the spread of parasite genetic adaptations; such as those conferring vaccine cross-protection against different strains affecting cattle and buffalo, or resistance to antiprotozoal drugs.

## References

Aktas, M., Dumanli, N., Angin, M., 2004. Cattle infestation by Hyalomma ticks and prevalence of Theileria in Hyalomma species in the east of Turkey. Vet Parasitol 119, 1–8.

Al-Hamidhi, S., M, H.T., Weir, W., Al-Fahdi, A., Johnson, E.H., Bobade, P., Alqamashoui, B., Beja-Pereira, A., Thompson, J., Kinnaird, J., Shiels, B., Tait, A., Babiker, H., 2015. Genetic Diversity and Population Structure of Theileria annulata in Oman. PLoS One 10, e0139581.

Al-Hosary, A., Abdel-Rady, A., Ahmed, L.S., Mohamed, A., 2010. Comparison between Using of BUPAQUONE ^®^ and Other Compounds in Treatment of Bovine Theileriosis. International Journal for Agro Veterinary and Medical Sciences 4, 3–7.

Bouattour, A., Darghouth, M.A., Ben Miled, L., 1996. Cattle infestation by Hyalomma ticks and prevalence of Theileria in H. detritum species in Tunisia. Vet Parasitol 65, 233–245.

Bouslikhane, M., M., K., H., O. 1998. La theilériose bovine au Maroc, investigations épidémiologiques et étude de l’impact sur la productivité des élevages. In: Résumé de communication orale, 15 ème congrès vétérinaire maghrébin.

Brown, C.G., 1990. Control of tropical theileriosis (Theileria annulata infection) of cattle. Parassitologia 32, 23–31.

Chatanga, E., Mosssad, E., Abdo Abubaker, H., Amin Alnour, S., Katakura, K., Nakao, R., Salim, B., 2019. Evidence of multiple point mutations in Theileria annulata cytochrome b gene incriminated in buparvaquone treatment failure. Acta Trop 191, 128–132.

Chaudhry, U., Ali, Q., Rashid, I., Shabbir, M.Z., Abbas, M., Numan, M., Evans, M., Ashraf, K., Morrison, I., Morrison, L., Sargison, N.D., 2019. Development of a deep amplicon sequencing method to determine the proportional species composition of piroplasm haemoprotozoa as an aid in their control. bioRxiv.

Darghouth, M.A., 2008. Review on the experience with live attenuated vaccines against tropical theileriosis in Tunisia: considerations for the present and implications for the future. Vaccine 26 Suppl 6, G4–G10.

Darghouth, M.A., Bouattour, A., Kilan, M., 1999. Tropical theileriosis in Tunisia: epidemiology and control. Parassitologia 41 Suppl 1, 33–36.

Estrada-Pena, A., Santos-Silva, M.M., 2005. The distribution of ticks (Acari: Ixodidae) of domestic livestock in Portugal. Experimental & applied acarology 36, 233–246.

Excoffier, L., Laval, G., Schneider, S., 2005. Arlequin (version 3.0): an integrated software package for population genetics data analysis. Evolutionary bioinformatics online 1, 47–50.

Excoffier, L., Smouse, P.E., Quattro, J.M., 1992. Analysis of molecular variance inferred from metric distances among DNA haplotypes: application to human mitochondrial DNA restriction data. Genetics 131, 479–491.

Gharbi, M., Darghouth, M.A., 2015. Control of tropical theileriosis (Theileria annulata infection in cattle) in North Africa. Asian Pacific Journal of Tropical Disease 5, 505–510.

Gharbi, M., Touay, A., Khayeche, M., Laarif, J., Jedidi, M., Sassi, L., Darghouth, M.A., 2011. Ranking control options for tropical theileriosis in at-risk dairy cattle in Tunisia, using benefit-cost analysis. Revue scientifique et technique (International Office of Epizootics) 30, 763–778.

Gomes, J., Salgueiro, P., Inacio, J., Amaro, A., Pinto, J., Tait, A., Shiels, B., Pereira da Fonseca, I., Santos-Gomes, G., Weir, W., 2016. Population diversity of Theileria annulata in Portugal. Infect Genet Evol 42, 14–19.

Guo, S.W., Thompson, E.A., 1992. Performing the exact test of Hardy-Weinberg proportion for multiple alleles. Biometrics 48, 361–372.

Jabbar, A., Abbas, T., Sandhu, Z.U., Saddiqi, H.A., Qamar, M.F., Gasser, R.B., 2015. Tick-borne diseases of bovines in Pakistan: major scope for future research and improved control. Parasites & vectors 8, 283.

Karim, S., Budachetri, K., Mukherjee, N., Williams, J., Kausar, A., Hassan, M.J., Adamson, S., Dowd, S.E., Apanskevich, D., Arijo, A., Sindhu, Z.U., Kakar, M.A., Khan, R.M.D., Ullah, S., Sajid, M.S., Ali, A., Iqbal, Z., 2017. A study of ticks and tick-borne livestock pathogens in Pakistan. PLoS neglected tropical diseases 11, e0005681.

Katzer, F., Carrington, M., Knight, P., Williamson, S., Tait, A., Morrison, I.W., Hall, R., 1994. Polymorphism of SPAG-1, a candidate antigen for inclusion in a sub-unit vaccine against Theileria annulata. Mol Biochem Parasitol 67, 1–10.

Kozich, J.J., Westcott, S.L., Baxter, N.T., Highlander, S.K., Schloss, P.D., 2013. Development of a dual-index sequencing strategy and curation pipeline for analyzing amplicon sequence data on the MiSeq Illumina sequencing platform. Appl. Environ. Microbiol. 79, 5112–5120.

Lawrence, J.A., 1979. The differential diagnosis of the bovine theilerias of Southern Africa. Journal of the South African Veterinary Association 50, 311–313.

Lefterova, M.I., Budvytiene, I., Sandlund, J., Farnert, A., Banaei, N., 2015. Simple Real-Time PCR and Amplicon Sequencing Method for Identification of Plasmodium Species in Human Whole Blood. J Clin Microbiol 53, 2251–2257.

Librado, P., Rozas, J., 2009. DnaSP v5: a software for comprehensive analysis of DNA polymorphism data. Bioinformatics 25, 1451–1452.

Mahmmod, Y.S., Elbalkemy, F.A., Klaas, I.C., Elmekkawy, M.F., Monazie, A.M., 2011. Clinical and haematological study on water buffaloes (Bubalus bubalis) and crossbred cattle naturally infected with Theileria annulata in Sharkia province, Egypt. Ticks Tick Borne Dis 2, 168–171.

Mhadhbi, M., Chaouch, M., Ajroud, K., Darghouth, M.A., BenAbderrazak, S., 2015. Sequence Polymorphism of Cytochrome b Gene in Theileria annulata Tunisian Isolates and Its Association with Buparvaquone Treatment Failure. PloS one 10, e0129678.

Mhadhbi, M., Naouach, A., Boumiza, A., Chaabani, M.F., BenAbderazzak, S., Darghouth, M.A., 2010. In vivo evidence for the resistance of Theileria annulata to buparvaquone. Vet Parasitol 169, 241–247.

Nourollahi-Fard, S.R., Khalili, M., Ghalekhani, N., 2015. Detection of Theileria annulata in blood samples of native cattle by PCR and smear method in Southeast of Iran. Journal of parasitic diseases : official organ of the Indian Society for Parasitology 39, 249–252.

Oura, C.A., Asiimwe, B.B., Weir, W., Lubega, G.W., Tait, A., 2005. Population genetic analysis and sub-structuring of Theileria parva in Uganda. Mol Biochem Parasitol 140, 229–239.

Oura, C.A., Tait, A., Asiimwe, B., Lubega, G.W., Weir, W., 2011. Haemoparasite prevalence and Theileria parva strain diversity in Cape buffalo (Syncerus caffer) in Uganda. Vet Parasitol 175, 212–219.

Pain, A., Renauld, H., Berriman, M., Murphy, L., Yeats, C.A., Weir, W., Kerhornou, A., Aslett, M., Bishop, R., Bouchier, C., Cochet, M., Coulson, R.M., Cronin, A., de Villiers, E.P., Fraser, A., Fosker, N., Gardner, M., Goble, A., Griffiths-Jones, S., Harris, D.E., Katzer, F., Larke, N., Lord, A., Maser, P., McKellar, S., Mooney, P., Morton, F., Nene, V., O’Neil, S., Price, C., Quail, M.A., Rabbinowitsch, E., Rawlings, N.D., Rutter, S., Saunders, D., Seeger, K., Shah, T., Squares, R., Squares, S., Tivey, A., Walker, A.R., Woodward, J., Dobbelaere, D.A., Langsley, G., Rajandream, M.A., McKeever, D., Shiels, B., Tait, A., Barrell, B., Hall, N., 2005. Genome of the host-cell transforming parasite Theileria annulata compared with T. parva. Science 309, 131–133.

Peakall, R., Smouse, P.E., 2012. GenAlEx 6.5: genetic analysis in Excel. Population genetic software for teaching and research--an update. Bioinformatics 28, 2537–2539.

Posada, D., 2008. jModelTest: phylogenetic model averaging. Molecular biology and evolution 25, 1253–1256.

Rice, W., 1989. Analyzing tables of statistical tests. Evolution 43.

Sargison, N.D., Shahzad, K., Mazeri, S., Chaudhry, U., 2019. A high throughput deep amplicon sequencing method to show the emergence and spread of Calicophoron daubneyi rumen fluke infection in United Kingdom cattle herds. Veterinary parasitology.

Sayin, F., Dincer, S., Karaer, Z., Cakmak, A., Inci, A., Yukari, B.A., Eren, H., Vatansever, Z., Nalbantoglu, S., 2003. Studies on the epidemiology of tropical theileriosis (Theileria annulata infection) in cattle in Central Anatolia, Turkey. Tropical animal health and production 35, 521–539.

Schloss, P.D., Westcott, S.L., Ryabin, T., Hall, J.R., Hartmann, M., Hollister, E.B., Lesniewski, R.A., Oakley, B.B., Parks, D.H., Robinson, C.J., 2009. Introducing mothur: open-source, platform-independent, community-supported software for describing and comparing microbial communities. Appl. Environ. Microbiol. 75, 7537–7541.

Schnittger, L., Katzer, F., Biermann, R., Shayan, P., Boguslawski, K., McKellar, S., Beyer, D., Shiels, B.R., Ahmed, J.S., 2002. Characterization of a polymorphic Theileria annulata surface protein (TaSP) closely related to PIM of Theileria parva: implications for use in diagnostic tests and subunit vaccines. Molecular and biochemical parasitology 120, 247–256.

Sharifiyazdi, H., Namazi, F., Oryan, A., Shahriari, R., Razavi, M., 2012. Point mutations in the Theileria annulata cytochrome b gene is associated with buparvaquone treatment failure. Vet Parasitol 187, 431–435.

Sivakumar, T., Hayashida, K., Sugimoto, C., Yokoyama, N., 2014. Evolution and genetic diversity of Theileria. Infect Genet Evol 27, 250–263.

Tait, A., Hall, F.R., 1990. Theileria annulata: control measures, diagnosis and the potential use of subunit vaccines. Rev Sci Tech 9, 387–403.

Uilenberg, G., Perie, N.M., Lawrence, J.A., de Vos, A.J., Paling, R.W., Spanjer, A.A., 1982. Causal agents of bovine theileriosis in southern Africa. Tropical animal health and production 14, 127–140.

Weir, W., Ben-Miled, L., Karagenc, T., Katzer, F., Darghouth, M., Shiels, B., Tait, A., 2007. Genetic exchange and sub-structuring in Theileria annulata populations. Mol Biochem Parasitol 154, 170–180.

Weir, W., Karagenc, T., Gharbi, M., Simuunza, M., Aypak, S., Aysul, N., Darghouth, M.A., Shiels, B., Tait, A., 2011. Population diversity and multiplicity of infection in Theileria annulata. Int J Parasitol 41, 193–203.

Yin, F., Liu, Z., Liu, J., Liu, A., Salih, D.A., Li, Y., Liu, G., Luo, J., Guan, G., Yin, H., 2018. Population Genetic Analysis of Theileria annulata from Six Geographical Regions in China, Determined on the Basis of Micro- and Mini-satellite Markers. Frontiers in genetics 9, 50.

Young, A.S., Brown, C.G., Burridge, M.J., Cunningham, M.P., Kirimi, I.M., Irvin, A.D., 1973. Observations on the cross-immunity between Theileria lawrencei (Serengeti) and Theileria parva (Muguga) in cattle. Int J Parasitol 3, 723–728.

