## Supplementary material for "Contrasting population genetics of cattle- and buffalo-derived *Theileria annulata* causing tropical theileriosis": Table 1

**Table 1:** Genetic comparison of *T. annulata* derived from buffalo and cattle based on six satellite and cytochrome b marker.

| **Satellite markers** | **Number of alleles (A_n_)** ^a^ | **Shared alleles (A_s_)** ^b^ | **Unique alleles (A_u_)** ^c^ |
| --- | --- | --- | --- |
| **Buffalo** |  |  |  |
| TS6 | 24 | 20 | 4 |
| TS8 | 26 | 26 | 0 |
| TS12 | 32 | 32 | 0 |
| TS16 | 23 | 20 | 3 |
| TS20 | 17 | 13 | 4 |
| TS31 | 20 | 20 | 0 |
| Mean | 23.7 | 20.7 | 1.8 |
| s.d. | 5.2 | 6.5 | 2.04 |
| **Cattle** |  |  |  |
| TS6 | 46 | 20 | 26 |
| TS8 | 37 | 26 | 11 |
| TS12 | 43 | 32 | 11 |
| TS16 | 28 | 20 | 9 |
| TS20 | 22 | 13 | 11 |
| TS31 | 32 | 20 | 12 |
| Mean | 34.7 | 20.7 | 13.3 |
| s.d. | 9.1 | 6.5 | 6.3 |
| **Mitochondrial marker** | **Number of alleles (A_n_)** ^a^ | **Shared alleles (A_s_)** ^b^ | **Unique alleles (A_u_)** ^c^ |
| **Buffalo** |  |  |  |
| Cytochrome b | 41 | 14 | 27 |
| **Cattle** |  |  |  |
| Cytochrome b | 79 | 14 | 65 |

^a^ Total number of alleles for buffalo and cattle derived *T. annulata*. ^b^ Total number of shared alleles. ^c^ Total number of unique alleles.
