## Supplementary material for "Contrasting population genetics of cattle- and buffalo-derived *Theileria annulata* causing tropical theileriosis": Table 2

**Table 2:** Genetic diversity data generated from genotyping of 18 buffalo- and 35 cattle- derived *T. annulata* populations using a panel of six satellite markers. Each value in the data set indicates the heterozygosity (He).

| **Buffalo** | **TS6** | **TS8** | **TS12** | **TS16** | **TS20** | **TS31** | **Mean** | **s.d.** | **Cattle** | **TS6** | **TS8** | **TS12** | **TS16** | **TS20** | **TS31** | **Mean** | **s.d.** |
| --- | --- | --- | --- | --- | --- | --- | --- | --- | --- | --- | --- | --- | --- | --- | --- | --- | --- |
| **CY7** | 0.857 | 0.964 | 0.929 | 0.892 | 0.929 | 0.929 | *0.935* | *0.047* | **CY2** | 0.909 | 0.926 | 0.996 | 0.952 | 0.887 | 0.948 | *0.937* | *0.038* |
| **CY16** | 0.867 | 0.933 | 0.933 | 0.933 | 0.867 | 0.933 | *0.911* | *0.034* | **CY6** | 0.858 | 0.968 | 0.995 | 0.932 | 0.916 | 0.953 | *0.937* | *0.048* |
| **CY92** | 0.821 | 0.929 | 0.964 | 0.857 | 0.857 | 0.893 | *0.887* | *0.053* | **CY79** | 0.945 | 0.989 | 0.967 | 0.912 | 0.890 | 0.967 | *0.945* | *0.037* |
| **CY109** | 0.909 | 0.545 | 0.939 | 0.939 | 0.818 | 0.909 | *0.843* | *0.153* | **CY82** | 0.954 | 0.993 | 0.961 | 0.954 | 0.895 | 0.895 | *0.942* | *0.039* |
| **CY157** | 0.857 | 0.857 | 0.963 | 0.964 | 0.929 | 0.929 | *0.923* | *0.057* | **CY89** | 0.792 | 0.992 | 0.992 | 0.933 | 0.942 | 0.967 | *0.936* | *0.075* |
| **CY103** | 0.857 | 0.857 | 0.964 | 0.786 | 0.929 | 0.857 | *0.875* | *0.063* | **CY84** | 0.912 | 0.868 | 0.978 | 0.956 | 0.791 | 0.945 | *0.908* | *0.069* |
| **CY15** | 0.911 | 0.800 | 0.911 | 0.911 | 0.800 | 0.978 | *0.885* | *0.071* | **CY20** | 0.929 | 0.973 | 0.893 | 0.964 | 0.893 | 0.857 | *0.923* | *0.053* |
| **CY98** | 0.802 | 0.945 | 0.812 | 0.857 | 0.934 | 0.934 | *0.912* | *0.071* | **CY90** | 0.882 | 0.993 | 0.980 | 0.850 | 0.791 | 0.935 | *0.905* | *0.079* |
| **CY129** | 0.545 | 0.939 | 0.825 | 0.939 | 0.970 | 0.894 | *0.881* | *0.168* | **CY26** | 0.909 | 0.993 | 0.980 | 0.869 | 0.856 | 0.922 | *0.922* | *0.056* |
| **CY142** | 0.857 | 0.571 | 0.464 | 0.908 | 0.714 | 0.821 | *0.738* | *0.196* | **CY27** | 0.893 | 0.964 | 0.879 | 0.821 | 0.857 | 0.964 | *0.917* | *0.070* |
| **CY141** | 0.800 | 0.822 | 0.733 | 0.911 | 0.822 | 0.822 | *0.819* | *0.057* | **CY28** | 0.956 | 0.978 | 0.956 | 0.956 | 0.978 | 0.933 | *0.959* | *0.017* |
| **CY138** | 0.848 | 0.939 | 0.985 | 0.909 | 0.939 | 0.894 | *0.919* | *0.047* | **CY30** | 0.842 | 0.950 | 0.992 | 0.825 | 0.808 | 0.950 | *0.894* | *0.078* |
| **CY148** | 0.689 | 0.911 | 0.876 | 0.983 | 0.956 | 0.933 | *0.915* | *0.116* | **CY34** | 0.983 | 0.965 | 0.893 | 0.922 | 0.909 | 0.957 | *0.956* | *0.035* |
| **CY126** | 0.935 | 0.933 | 0.933 | 0.849 | 0.933 | 0.933 | *0.956* | *0.034* | **CY39** | 0.989 | 0.967 | 0.956 | 0.934 | 0.945 | 0.901 | *0.956* | *0.030* |
| **CY149** | 0.571 | 0.857 | 0.929 | 0.943 | 0.893 | 0.857 | *0.851* | *0.147* | **CY38** | 0.857 | 0.929 | 0.929 | 0.964 | 0.857 | 0.857 | *0.949* | *0.047* |
| **CY139** | 0.933 | 0.733 | 0.933 | 0.893 | 0.867 | 0.867 | *0.889* | *0.091* | **CY40** | 0.912 | 0.956 | 0.989 | 0.857 | 0.846 | 0.912 | *0.899* | *0.055* |
| **CY123** | 0.739 | 0.941 | 0.993 | 0.804 | 0.869 | 0.941 | *0.881* | *0.096* | **CY41** | 0.867 | 0.978 | 0.956 | 0.822 | 0.822 | 0.911 | *0.912* | *0.067* |
| **CY133** | 0.864 | 0.939 | 0.985 | 0.810 | 0.924 | 0.864 | *0.899* | *0.061* | **CY37** | 0.857 | 0.964 | 0.964 | 0.857 | 0.964 | 0.929 | *0.893* | *0.053* |
|  |  |  |  |  |  |  | *0.874* | *0.028* | **CY160** | 0.939 | 0.993 | 0.970 | 0.939 | 0.970 | 0.970 | *0.923* | *0.023* |
|  |  |  |  |  |  |  |  |  | **CY162** | 0.864 | 0.970 | 0.985 | 0.864 | 0.924 | 0.818 | *0.965* | *0.066* |
|  |  |  |  |  |  |  |  |  | **CY154** | 0.893 | 0.964 | 0.929 | 0.857 | 0.857 | 0.893 | *0.904* | *0.042* |
|  |  |  |  |  |  |  |  |  | **CY131** | 0.867 | 0.978 | 0.889 | 0.889 | 0.911 | 0.911 | *0.899* | *0.038* |
|  |  |  |  |  |  |  |  |  | **CY130** | 0.924 | 0.985 | 0.985 | 0.909 | 0.924 | 0.909 | *0.907* | *0.036* |
|  |  |  |  |  |  |  |  |  | **CY120** | 0.934 | 0.912 | 0.890 | 0.802 | 0.912 | 0.846 | *0.939* | *0.069* |
|  |  |  |  |  |  |  |  |  | **CY119** | 0.899 | 0.857 | 0.912 | 0.857 | 0.964 | 0.964 | *0.897* | *0.067* |
|  |  |  |  |  |  |  |  |  | **CY134** | 0.943 | 0.956 | 0.978 | 0.923 | 0.934 | 0.945 | *0.940* | *0.029* |
|  |  |  |  |  |  |  |  |  | **CY150** | 0.933 | 0.933 | 0.933 | 0.933 | 0.933 | 0.933 | *0.956* | *0.000* |
|  |  |  |  |  |  |  |  |  | **CY173** | 0.600 | 0.821 | 0.867 | 0.733 | 0.733 | 0.733 | *0.933* | *0.140* |
|  |  |  |  |  |  |  |  |  | **CY124** | 0.800 | 0.812 | 0.923 | 0.867 | 0.867 | 0.978 | *0.800* | *0.082* |
|  |  |  |  |  |  |  |  |  | **CY175** | 0.945 | 0.923 | 0.967 | 0.890 | 0.890 | 0.978 | *0.926* | *0.040* |
|  |  |  |  |  |  |  |  |  | **CY177** | 0.939 | 0.985 | 0.970 | 0.894 | 0.894 | 0.955 | *0.951* | *0.031* |
|  |  |  |  |  |  |  |  |  | **CY136** | 0.886 | 0.994 | 0.882 | 0.902 | 0.902 | 0.938 | *0.947* | *0.050* |
|  |  |  |  |  |  |  |  |  | **CY151** | 0.887 | 0.957 | 0.991 | 0.870 | 0.870 | 0.935 | *0.936* | *0.121* |
|  |  |  |  |  |  |  |  |  | **CY145** | 0.870 | 0.974 | 0.974 | 0.883 | 0.883 | 0.948 | *0.882* | *0.061* |
|  |  |  |  |  |  |  |  |  | **CY33** | 0.963 | 0.995 | 0.995 | 0.942 | 0.942 | 0.921 | *0.913* | *0.128* |
|  |  |  |  |  |  |  |  |  |  |  |  |  |  |  |  | *0.912* | *0.056* |

TS; satellite markers across all populations, CY; total number of *T. annulata* populations in buffalo and cattle, Mean and standard deviation (s.d.) demonstrate the heterozygosity (H_e_) in each population for six markers.
