## Supplementary material for "Contrasting population genetics of cattle- and buffalo-derived *Theileria annulata* causing tropical theileriosis": Table 3

**Table 3.** Summary of the genetic diversity data for the cytochrome b locus of 31 buffalo- and 54 cattle-derived *T. annulate* populations.

| **Buffalo** | **He** | | **S** | | **π** | **S_θ_** | | **k** | | **Cattle** | **He** | **S** | **π** | **S_θ_** | **k** |
| --- | --- | --- | --- | --- | --- | --- | --- | --- | --- | --- | --- | --- | --- | --- | --- |
| **CY91** | 0.869 | | 12 | | 0.00489 | 0.00587 | | 2.524 | | **CY1** | 0.571 | 3 | 0.00332 | 0.00224 | 1.714 |
| **CY92** | 0.556 | | 2 | | 0.00215 | 0.00137 | | 1.111 | | **CY2** | 0.800 | 6 | 0.00517 | 0.00350 | 2.667 |
| **CY94** | N/A | |  | |  |  | |  | | **CY3** | 0.556 | 3 | 0.00323 | 0.00206 | 1.667 |
| **CY96** | 0.545 | | 3 | | 0.00317 | 0.00193 | | 1.636 | | **CY6** | 0.789 | 7 | 0.00535 | 0.00382 | 2.763 |
| **CY98** | 0.870 | | 10 | | 0.00629 | 0.00519 | | 3.246 | | **CY20** | 0.870 | 10 | 0.00640 | 0.00519 | 3.304 |
| **CY103** | N/A | |  | |  |  | |  | | **CY25** | 0.556 | 3 | 0.00323 | 0.00206 | 1.667 |
| **CY105** | N/A | |  | |  |  | |  | | **CY26** | 0.842 | 8 | 0.00588 | 0.00437 | 3.032 |
| **CY107** | 0.571 | | 3 | | 0.00332 | 0.00224 | | 1.714 | | **CY27** | 0.833 | 8 | 0.00581 | 0.00411 | 3.000 |
| **CY109** | 0.800 | | 7 | | 0.00568 | 0.00409 | | 2.933 | | **CY28** | 0.800 | 7 | 0.00543 | 0.00409 | 2.800 |
| **CY110** | 0.800 | | 9 | | 0.00724 | 0.00526 | | 3.733 | | **CY29** | 0.556 | 4 | 0.00431 | 0.00274 | 2.222 |
| **CY111** | N/A | |  | |  |  | |  | | **CY30** | 0.714 | 5 | 0.00461 | 0.00298 | 2.381 |
| **CY112** | N/A | |  | |  |  | |  | | **CY31** | 0.545 | 3 | 0.00317 | 0.00193 | 1.636 |
| **CY113** | 0.571 | | 3 | | 0.00332 | 0.00224 | | 1.714 | | **CY32** | 0.789 | 7 | 0.00586 | 0.00382 | 3.026 |
| **CY117** | 0.727 | | 5 | | 0.00470 | 0.00321 | | 2.424 | | **CY33** | 0.789 | 6 | 0.00510 | 0.00328 | 2.632 |
| **CY123** | 0.842 | | 10 | | 0.00718 | 0.00546 | | 3.705 | | **CY34** | 0.556 | 4 | 0.00431 | 0.00274 | 2.222 |
| **CY126** | 0.545 | | 3 | | 0.00317 | 0.00193 | | 1.636 | | **CY35** | 0.706 | 4 | 0.00365 | 0.00225 | 1.882 |
| **CY128** | 0.897 | | 11 | | 0.00603 | 0.00501 | | 3.109 | | **CY36** | 0.545 | 3 | 0.00317 | 0.00193 | 1.636 |
| **CY129** | 0.833 | | 7 | | 0.00452 | 0.00359 | | 2.333 | | **CY37** | 0.789 | 5 | 0.00433 | 0.00273 | 2.237 |
| **CY133** | 0.842 | | 7 | | 0.00522 | 0.00382 | | 2.695 | | **CY38** | 0.789 | 7 | 0.00535 | 0.00382 | 2.763 |
| **CY138** | 0.889 | | 10 | | 0.00558 | 0.00498 | | 2.878 | | **CY39** | 0.913 | 8 | 0.00651 | 0.00415 | 3.359 |
| **CY139** | N/A | |  | |  |  | |  | | **CY40** | 0.842 | 9 | 0.00685 | 0.00492 | 3.537 |
| **CY141** | N/A | |  | |  |  | |  | | **CY41** | 0.789 | 5 | 0.00433 | 0.00273 | 2.237 |
| **CY142** | N/A | |  | |  |  | |  | | **CY42** | 0.862 | 9 | 0.00613 | 0.00440 | 3.161 |
| **CY148** | 0.727 | | 4 | | 0.00376 | 0.00257 | | 1.939 | | **CY79** | 0.897 | 13 | 0.00658 | 0.00592 | 3.397 |
| **CY149** | 0.571 | | 3 | | 0.00332 | 0.00224 | | 1.714 | | **CY80** | N/A |  |  |  |  |
| **CY157** | 0.556 | | 3 | | 0.00323 | 0.00206 | | 1.667 | | **CY82** | 0.789 | 7 | 0.00561 | 0.00382 | 2.895 |
| **CY7** | 0.556 | | 3 | | 0.00323 | 0.00206 | | 1.667 | | **CY83** | 0.800 | 7 | 0.00568 | 0.00409 | 2.933 |
| **CY8** | 0.714 | | 5 | | 0.00461 | 0.00298 | | 2.381 | | **CY84** | 0.882 | 10 | 0.00604 | 0.00563 | 3.118 |
| **CY15** | 0.556 | | 3 | | 0.00323 | 0.00206 | | 1.667 | | **CY86** | 0.857 | 8 | 0.00598 | 0.00477 | 3.086 |
| **CY16** | 0.545 | | 3 | | 0.00317 | 0.00193 | | 1.636 | | **CY87** | 0.842 | 9 | 0.00588 | 0.00492 | 3.032 |
| **CY19** | 0.900 | | 10 | | 0.00581 | 0.00539 | | 3.000 | | **CY88** | 0.714 | 5 | 0.00461 | 0.00298 | 2.381 |
| **Total** | 0.861 | | 47 | | 0.00526 | 0.01654 | | 2.712 | | **CY89** | 0.870 | 9 | 0.00573 | 0.00467 | 2.957 |
|  |  |  | |  | | |  | |  | **CY90** | 0.974 | 21 | 0.00710 | 0.01002 | 3.662 |
|  |  |  | |  | | |  | |  | **CY119** | 0.800 | 7 | 0.00543 | 0.00409 | 2.800 |
|  |  |  | |  | | |  | |  | **CY120** | 0.842 | 8 | 0.00522 | 0.00437 | 2.695 |
|  |  |  | |  | | |  | |  | **CY124** | N/A |  |  |  |  |
|  |  |  | |  | | |  | |  | **CY130** | 0.800 | 6 | 0.00517 | 0.00350 | 2.667 |
|  |  |  | |  | | |  | |  | **CY131** | 0.857 | 8 | 0.00598 | 0.00477 | 3.086 |
|  |  |  | |  | | |  | |  | **CY134** | 0.727 | 5 | 0.00470 | 0.00321 | 2.424 |
|  |  |  | |  | | |  | |  | **CY135** | 0.727 | 5 | 0.00470 | 0.00321 | 2.424 |
|  |  |  | |  | | |  | |  | **CY136** | 0.923 | 11 | 0.00596 | 0.00670 | 3.077 |
|  |  |  | |  | | |  | |  | **CY140** | 0.727 | 5 | 0.00470 | 0.00321 | 2.424 |
|  |  |  | |  | | |  | |  | **CY145** | 0.889 | 11 | 0.00656 | 0.00548 | 3.386 |
|  |  |  | |  | | |  | |  | **CY150** | 0.862 | 9 | 0.00601 | 0.00440 | 3.103 |
|  |  |  | |  | | |  | |  | **CY151** | N/A |  |  |  |  |
|  |  |  | |  | | |  | |  | **CY153** | N/A |  |  |  |  |
|  |  |  | |  | | |  | |  | **CY154** | N/A |  |  |  |  |
|  |  |  | |  | | |  | |  | **CY158** | N/A |  |  |  |  |
|  |  |  | |  | | |  | |  | **CY160** | N/A |  |  |  |  |
|  |  |  | |  | | |  | |  | **CY162** | 0.727 | 6 | 0.00564 | 0.00385 | 2.909 |
|  |  |  | |  | | |  | |  | **CY163** | N/A |  |  |  |  |
|  |  |  | |  | | |  | |  | **CY173** | N/A |  |  |  |  |
|  |  |  | |  | | |  | |  | **CY175** | N/A |  |  |  |  |
|  |  |  | |  | | |  | |  | **CY177** | 0.727 | 5 | 0.00470 | 0.00321 | 2.424 |
|  |  |  | |  | | |  | |  | **Total** | 0.931 | 49 | 0.00615 | 0.05188 | 3.174 |

Total number of *T. annulata* populations in buffalo and cattle (CY), heterozygosity (He), the number of segregating sites (S), nucleotide diversity (π), the mean number of pairwise differences (k), the mutation parameter based on an infinite site equilibrium model, and the mutations parameter based on segregating sites (Sθ).
