## Supplementary material for "Contrasting population genetics of cattle- and buffalo-derived *Theileria annulata* causing tropical theileriosis": Table 4

**Table 4:** Multilocus genotype data of 18 buffalo- and 35 cattle-derived *T. annulata* populations, based on a panel of six satellite markers. Each value in the data set indicates the numbers of genotypes per population.

| **Buffalo** | **TS6** | **TS8** | **TS12** | **TS16** | **TS20** | **TS31** | **Mean** | **s.d.** | **Cattle** | **TS6** | **TS8** | **TS12** | **TS16** | **TS20** | **TS31** | **Mean** | **s.d.** |
| --- | --- | --- | --- | --- | --- | --- | --- | --- | --- | --- | --- | --- | --- | --- | --- | --- | --- |
| **CY7** | 4 | 6 | 6 | 8 | 6 | 6 | *6* | *1.26* | **CY2** | 8 | 10 | 21 | 12 | 8 | 12 | *11.83* | *4.83* |
| **CY16** | 4 | 5 | 5 | 5 | 4 | 4 | *4.5* | *0.55* | **CY6** | 6 | 13 | 19 | 10 | 7 | 11 | *11* | *4.69* |
| **CY92** | 4 | 6 | 7 | 5 | 5 | 5 | *5.33* | *1.03* | **CY79** | 9 | 13 | 11 | 7 | 6 | 11 | *9.5* | *2.66* |
| **CY109** | 7 | 2 | 9 | 8 | 5 | 7 | *6.33* | *2.5* | **CY82** | 4 | 17 | 11 | 11 | 8 | 7 | *9.5* | *4.55* |
| **CY157** | 4 | 4 | 8 | 7 | 6 | 6 | *5.83* | *1.6* | **CY89** | 7 | 15 | 15 | 9 | 10 | 12 | *11.33* | *3.27* |
| **CY103** | 4 | 5 | 7 | 4 | 6 | 4 | *5* | *1.26* | **CY84** | 7 | 6 | 12 | 6 | 4 | 10 | *7.5* | *2.95* |
| **CY15** | 6 | 4 | 6 | 6 | 4 | 9 | *5.83* | *1.83* | **CY20** | 7 | 8 | 5 | 7 | 4 | 4 | *5.83* | *1.72* |
| **CY98** | 4 | 9 | 14 | 6 | 9 | 8 | *8.33* | *3.39* | **CY90** | 8 | 17 | 15 | 6 | 4 | 9 | *9.83* | *5.12* |
| **CY129** | 2 | 8 | 12 | 9 | 10 | 6 | *7.83* | *3.49* | **CY26** | 8 | 17 | 15 | 6 | 6 | 9 | *10.17* | *4.71* |
| **CY142** | 4 | 2 | 3 | 8 | 3 | 4 | *4* | *2.1* | **CY27** | 5 | 7 | 8 | 4 | 5 | 7 | *6* | *1.55* |
| **CY141** | 4 | 4 | 3 | 6 | 4 | 4 | *4.17* | *0.98* | **CY28** | 8 | 9 | 8 | 8 | 9 | 7 | *8.17* | *0.75* |
| **CY138** | 5 | 7 | 10 | 7 | 8 | 6 | *7.17* | *1.72* | **CY30** | 5 | 10 | 15 | 5 | 5 | 10 | *8.33* | *4.08* |
| **CY148** | 3 | 5 | 10 | 10 | 8 | 7 | *7.17* | *2.79* | **CY34** | 18 | 15 | 22 | 9 | 8 | 12 | *14* | *5.4* |
| **CY126** | 6 | 5 | 5 | 6 | 5 | 5 | *5.33* | *0.52* | **CY39** | 13 | 10 | 10 | 8 | 9 | 7 | *9.5* | *2.07* |
| **CY149** | 2 | 4 | 6 | 8 | 5 | 5 | *5* | *2* | **CY38** | 5 | 6 | 6 | 7 | 4 | 4 | *5.33* | *1.21* |
| **CY139** | 5 | 3 | 5 | 6 | 4 | 4 | *4.5* | *1.05* | **CY40** | 7 | 10 | 13 | 5 | 6 | 7 | *8* | *2.97* |
| **CY123** | 4 | 10 | 17 | 5 | 6 | 10 | *8.67* | *4.8* | **CY41** | 5 | 8 | 8 | 4 | 4 | 6 | *5.83* | *1.83* |
| **CY133** | 5 | 8 | 11 | 4 | 5 | 5 | *6.33* | *2.66* | **CY37** | 5 | 7 | 7 | 5 | 7 | 6 | *6.17* | *0.98* |
|  |  |  |  |  |  |  | *6.25* | *1.68* | **CY160** | 8 | 12 | 10 | 8 | 10 | 10 | *9.67* | *1.51* |
|  |  |  |  |  |  |  |  |  | **CY162** | 5 | 10 | 11 | 5 | 7 | 4 | *7* | *2.9* |
|  |  |  |  |  |  |  |  |  | **CY154** | 5 | 7 | 6 | 5 | 5 | 5 | *5.5* | *0.84* |
|  |  |  |  |  |  |  |  |  | **CY131** | 5 | 9 | 5 | 5 | 6 | 6 | *6* | *1.55* |
|  |  |  |  |  |  |  |  |  | **CY130** | 6 | 11 | 11 | 7 | 7 | 6 | *8* | *2.37* |
|  |  |  |  |  |  |  |  |  | **CY120** | 8 | 14 | 6 | 4 | 7 | 5 | *7.33* | *3.56* |
|  |  |  |  |  |  |  |  |  | **CY119** | 8 | 4 | 8 | 5 | 7 | 7 | *6.5* | *1.64* |
|  |  |  |  |  |  |  |  |  | **CY134** | 14 | 10 | 12 | 8 | 8 | 9 | *10.17* | *2.4* |
|  |  |  |  |  |  |  |  |  | **CY150** | 5 | 5 | 5 | 5 | 5 | 5 | *5* | *0* |
|  |  |  |  |  |  |  |  |  | **CY173** | 2 | 6 | 4 | 3 | 4 | 3 | *3.67* | *1.37* |
|  |  |  |  |  |  |  |  |  | **CY124** | 4 | 10 | 10 | 5 | 6 | 9 | *7.33* | *2.66* |
|  |  |  |  |  |  |  |  |  | **CY175** | 8 | 14 | 11 | 6 | 8 | 12 | *9.83* | *2.99* |
|  |  |  |  |  |  |  |  |  | **CY177** | 8 | 11 | 10 | 6 | 8 | 9 | *8.67* | *1.75* |
|  |  |  |  |  |  |  |  |  | **CY136** | 7 | 24 | 26 | 8 | 8 | 11 | *14* | *8.65* |
|  |  |  |  |  |  |  |  |  | **CY151** | 7 | 14 | 20 | 6 | 3 | 10 | *10* | *6.16* |
|  |  |  |  |  |  |  |  |  | **CY145** | 6 | 15 | 16 | 7 | 5 | 11 | *10* | *4.73* |
|  |  |  |  |  |  |  |  |  | **CY33** | 13 | 19 | 19 | 10 | 3 | 9 | *12.17* | *6.21* |
|  |  |  |  |  |  |  |  |  |  |  |  |  |  |  |  | *8.60* | *2.52* |

TS; satellite markers across all populations, CY; total number of *T. annulata* populations in buffalo and cattle, Mean and standard deviation (s.d.) demonstrate the genotypes in each population for six markers.
