## Supplementary Table S1 for "Contrasting population genetics of cattle- and buffalo-derived *Theileria annulata* causing tropical theileriosis"

**Supplementary Table S1.** Primer sequences for Illumina MiSeq Library preparation. Tann-ForAdp/Tann-RevAdp primer sequence are underlined, N’s are bolded.

| **Sequences (5'-3')** | **Primer Name** |
| --- | --- |
| TCGTCGGCAGCGTCAGATGTGTATAAGAGACAGAAGTATAGCAACTGCTTTTGTT | Tann-ForAdp |
| TCGTCGGCAGCGTCAGATGTGTATAAGAGACAG**N**AAGTATAGCAACTGCTTTTGTT | Tann-ForAdp1N |
| TCGTCGGCAGCGTCAGATGTGTATAAGAGACAG**NN**AAGTATAGCAACTGCTTTTGTT | Tann-ForAdp2N |
| TCGTCGGCAGCGTCAGATGTGTATAAGAGACAG**NNN**AAGTATAGCAACTGCTTTTGTT | Tann-ForRdp3N |
| GTCTCGTGGGCTCGGAGATGTGTATAAGAGACAGTCCTGCCATTGCCAAAAGTC | Tann-RevAdp |
| GTCTCGTGGGCTCGGAGATGTGTATAAGAGACAG**N**TCCTGCCATTGCCAAAAGTC | Tann-RevAdp1N |
| GTCTCGTGGGCTCGGAGATGTGTATAAGAGACAG**NN**TCCTGCCATTGCCAAAAGTC | Tann-RevAdp2N |
| GTCTCGTGGGCTCGGAGATGTGTATAAGAGACAG**NNN**TCCTGCCATTGCCAAAAGTC | Tann-Revdp3N |
