## Supplementary Table S3 for "Contrasting population genetics of cattle- and buffalo-derived *Theileria annulata* causing tropical theileriosis"

| **Name** | **Chromosome** | **Location of repeated region** | **Consensus repeat sequence** | ***T. annulata* size range** | **PCR primers 5′–3′** | **Annealing temp. (°C)** |
| --- | --- | --- | --- | --- | --- | --- |
| TS6 | 4 | Intron | TAATTATAGG | 301–466 | F catcctttgacctactgattgtac | 60 |
|  |  |  |  |  | R cggtagtaccagttaatactgtc |  |
| TS8 | 3 | Exon | TATTATTTAATG | 195–356 | F taaacgattaaaatcaagtg | 55 |
|  |  |  |  |  | R attggaaatggtgaaataatgag |  |
| TS12 | 3 | Exon | AATACT | 237–376 | F gatgatagaggaattgatatgac | 55 |
|  |  |  |  |  | R ggaaatatcacaattaagattc |  |
| TS16 | 1 | Intergenic | TAA | 345–439 | F ccaatgtcaacagtatgatg | 55 |
|  |  |  |  |  | R gagtaagaagtaccactactg |  |
| TS20 | 2 | Intron | ATTATTACTA | 187–310 | F ccttcatgatctacatctgatgc | 60 |
|  |  |  |  |  | R ggctgaatgggtacctgttc |  |
| TS31 | 2 | Non-coding region | AATTTATCCTGAATTATAGA | 203–385 | F gttatcttcttgctattatagc | 50 |
|  |  |  |  |  | R gtattaaaatctataagattc |  |

**Supplementary Table S3.** Sequences and allele ranges for two microsatellite (TS12, TS16) and four minisatellite markers (TS6, TS8, TS20, TS31 Sequences.
