## Supplementary Table S4 for "Contrasting population genetics of cattle- and buffalo-derived *Theileria annulata* causing tropical theileriosis"

**Supplementary Table S4** The values of fixation index (F_ST_) based on genotyping 35 cattle and 18 buffalo derived *T. annulata* populations based on the panel of six satellite markers.

| **Cattle** | **CY2** | **CY6** | **CY79** | **CY82** | **CY89** | **CY84** | **CY20** | **CY90** | **CY26** | **CY27** | **CY28** | **CY30** | **CY34** | **CY39** | **CY38** | **CY40** | **CY41** | **CY37** | **CY160** | **CY162** | **CY154** | **CY131** | **CY130** | **CY120** | **CY119** | **CY134** | **CY150** | **CY173** | **CY124** | **CY175** | **CY177** | **CY136** | **CY151** | **CY145** |
| --- | --- | --- | --- | --- | --- | --- | --- | --- | --- | --- | --- | --- | --- | --- | --- | --- | --- | --- | --- | --- | --- | --- | --- | --- | --- | --- | --- | --- | --- | --- | --- | --- | --- | --- |
| **CY6** | 0.010 |  |  |  |  |  |  |  |  |  |  |  |  |  |  |  |  |  |  |  |  |  |  |  |  |  |  |  |  |  |  |  |  |  |
| **CY79** | 0.015 | 0.008 |  |  |  |  |  |  |  |  |  |  |  |  |  |  |  |  |  |  |  |  |  |  |  |  |  |  |  |  |  |  |  |  |
| **CY82** | 0.010 | 0.011 | 0.015 |  |  |  |  |  |  |  |  |  |  |  |  |  |  |  |  |  |  |  |  |  |  |  |  |  |  |  |  |  |  |  |
| **CY89** | 0.012 | 0.020 | 0.007 | 0.002 |  |  |  |  |  |  |  |  |  |  |  |  |  |  |  |  |  |  |  |  |  |  |  |  |  |  |  |  |  |  |
| **CY84** | 0.019 | 0.021 | 0.015 | 0.001 | 0.012 |  |  |  |  |  |  |  |  |  |  |  |  |  |  |  |  |  |  |  |  |  |  |  |  |  |  |  |  |  |
| **CY20** | -0.004 | 0.007 | 0.013 | 0.021 | 0.024 | 0.029 |  |  |  |  |  |  |  |  |  |  |  |  |  |  |  |  |  |  |  |  |  |  |  |  |  |  |  |  |
| **CY90** | 0.035 | 0.037 | 0.036 | 0.025 | 0.035 | 0.039 | 0.035 |  |  |  |  |  |  |  |  |  |  |  |  |  |  |  |  |  |  |  |  |  |  |  |  |  |  |  |
| **CY26** | 0.018 | 0.025 | 0.018 | 0.018 | 0.018 | 0.025 | 0.016 | -0.028 |  |  |  |  |  |  |  |  |  |  |  |  |  |  |  |  |  |  |  |  |  |  |  |  |  |  |
| **CY27** | 0.019 | 0.032 | 0.017 | 0.020 | 0.011 | 0.032 | 0.024 | 0.043 | 0.031 |  |  |  |  |  |  |  |  |  |  |  |  |  |  |  |  |  |  |  |  |  |  |  |  |  |
| **CY28** | 0.006 | 0.006 | 0.003 | -0.005 | 0.009 | 0.017 | 0.015 | 0.014 | -0.001 | 0.021 |  |  |  |  |  |  |  |  |  |  |  |  |  |  |  |  |  |  |  |  |  |  |  |  |
| **CY30** | 0.022 | 0.014 | 0.032 | 0.022 | 0.034 | 0.017 | 0.022 | 0.058 | 0.028 | 0.057 | 0.025 |  |  |  |  |  |  |  |  |  |  |  |  |  |  |  |  |  |  |  |  |  |  |  |
| **CY34** | 0.006 | 0.006 | -0.006 | -0.004 | 0.001 | 0.010 | 0.011 | 0.017 | 0.006 | 0.022 | -0.010 | 0.024 |  |  |  |  |  |  |  |  |  |  |  |  |  |  |  |  |  |  |  |  |  |  |
| **CY39** | 0.006 | 0.013 | -0.048 | -0.016 | 0.007 | -0.001 | 0.021 | 0.022 | 0.013 | 0.010 | 0.005 | 0.017 | -0.006 |  |  |  |  |  |  |  |  |  |  |  |  |  |  |  |  |  |  |  |  |  |
| **CY38** | 0.041 | 0.033 | 0.020 | 0.025 | 0.029 | 0.045 | 0.026 | 0.060 | 0.043 | 0.040 | 0.028 | 0.046 | 0.017 | 0.028 |  |  |  |  |  |  |  |  |  |  |  |  |  |  |  |  |  |  |  |  |
| **CY40** | 0.029 | 0.012 | 0.019 | 0.021 | 0.009 | 0.006 | 0.026 | 0.049 | 0.037 | 0.014 | 0.015 | 0.040 | 0.014 | 0.010 | 0.029 |  |  |  |  |  |  |  |  |  |  |  |  |  |  |  |  |  |  |  |
| **CY41** | 0.051 | 0.043 | 0.048 | 0.026 | 0.048 | 0.055 | 0.045 | 0.036 | 0.055 | 0.033 | 0.039 | 0.064 | 0.028 | 0.041 | 0.036 | 0.030 |  |  |  |  |  |  |  |  |  |  |  |  |  |  |  |  |  |  |
| **CY37** | 0.019 | 0.009 | 0.011 | 0.017 | 0.024 | 0.029 | 0.013 | 0.040 | 0.030 | 0.040 | 0.015 | 0.039 | 0.004 | 0.011 | 0.039 | 0.051 | 0.041 |  |  |  |  |  |  |  |  |  |  |  |  |  |  |  |  |  |
| **CY160** | 0.001 | -0.005 | -0.004 | -0.001 | 0.005 | 0.009 | -0.004 | 0.010 | 0.005 | 0.010 | -0.007 | 0.009 | -0.007 | 0.001 | 0.010 | 0.015 | -0.018 | -0.042 |  |  |  |  |  |  |  |  |  |  |  |  |  |  |  |  |
| **CY162** | 0.020 | 0.018 | 0.021 | 0.015 | 0.024 | 0.026 | 0.023 | 0.037 | 0.022 | 0.049 | 0.017 | 0.035 | 0.013 | 0.015 | 0.035 | 0.037 | 0.043 | -0.009 | -0.003 |  |  |  |  |  |  |  |  |  |  |  |  |  |  |  |
| **CY154** | 0.021 | 0.027 | 0.023 | 0.021 | 0.028 | 0.042 | 0.029 | 0.026 | 0.025 | 0.058 | 0.013 | 0.058 | 0.000 | 0.016 | 0.033 | 0.035 | 0.032 | 0.020 | 0.005 | -0.032 |  |  |  |  |  |  |  |  |  |  |  |  |  |  |
| **CY131** | 0.031 | 0.025 | 0.035 | 0.024 | 0.028 | 0.038 | 0.011 | 0.035 | 0.028 | 0.044 | 0.004 | 0.049 | 0.007 | 0.026 | 0.026 | 0.015 | 0.024 | 0.007 | -0.014 | 0.004 | 0.024 |  |  |  |  |  |  |  |  |  |  |  |  |  |
| **CY130** | 0.021 | 0.003 | 0.013 | 0.007 | 0.013 | 0.014 | 0.019 | 0.037 | 0.026 | 0.040 | -0.006 | 0.038 | 0.005 | 0.018 | 0.009 | 0.019 | 0.008 | 0.012 | -0.021 | 0.016 | 0.002 | -0.011 |  |  |  |  |  |  |  |  |  |  |  |  |
| **CY120** | 0.035 | 0.030 | 0.027 | 0.021 | 0.026 | 0.039 | 0.040 | 0.027 | 0.053 | 0.021 | 0.045 | 0.018 | 0.016 | 0.030 | 0.020 | 0.027 | 0.035 | 0.003 | 0.033 | 0.033 | 0.011 | 0.020 | -0.002 |  |  |  |  |  |  |  |  |  |  |  |
| **CY119** | 0.004 | 0.017 | 0.012 | 0.002 | 0.008 | 0.017 | 0.015 | 0.027 | 0.023 | 0.001 | -0.007 | 0.048 | -0.004 | 0.001 | 0.005 | 0.013 | 0.001 | -0.005 | -0.011 | -0.011 | -0.030 | -0.005 | -0.023 | 0.007 |  |  |  |  |  |  |  |  |  |  |
| **CY134** | 0.015 | 0.011 | 0.006 | 0.003 | 0.009 | 0.025 | 0.017 | 0.022 | 0.018 | 0.019 | -0.007 | 0.036 | -0.004 | 0.000 | 0.012 | 0.006 | 0.005 | 0.005 | -0.010 | 0.008 | -0.007 | -0.004 | -0.024 | -0.017 | -0.055 |  |  |  |  |  |  |  |  |  |
| **CY150** | 0.009 | 0.011 | 0.011 | 0.016 | 0.016 | 0.035 | 0.028 | 0.016 | 0.014 | 0.039 | -0.003 | 0.051 | -0.010 | 0.009 | 0.013 | 0.018 | 0.015 | 0.010 | -0.009 | -0.040 | -0.132 | 0.012 | -0.019 | 0.003 | -0.050 | -0.023 |  |  |  |  |  |  |  |  |
| **CY173** | 0.076 | 0.081 | 0.076 | 0.081 | 0.087 | 0.094 | 0.078 | 0.087 | 0.092 | 0.101 | 0.051 | 0.086 | 0.074 | 0.072 | 0.107 | 0.088 | 0.110 | 0.098 | 0.059 | 0.074 | 0.084 | 0.114 | 0.086 | 0.098 | 0.091 | 0.085 | 0.069 |  |  |  |  |  |  |  |
| **CY124** | 0.011 | 0.018 | 0.000 | 0.011 | 0.015 | 0.011 | 0.007 | 0.019 | 0.017 | 0.019 | 0.013 | 0.038 | 0.013 | 0.007 | 0.039 | 0.026 | 0.034 | -0.037 | -0.021 | -0.003 | 0.020 | -0.013 | 0.014 | 0.031 | -0.006 | 0.007 | 0.010 | 0.088 |  |  |  |  |  |  |
| **CY175** | 0.018 | 0.011 | 0.005 | -0.002 | 0.020 | 0.010 | 0.008 | 0.015 | 0.010 | 0.028 | -0.004 | 0.012 | 0.000 | -0.016 | 0.018 | 0.012 | 0.019 | -0.005 | -0.021 | 0.020 | 0.035 | 0.005 | 0.008 | 0.024 | 0.013 | 0.002 | 0.022 | 0.062 | 0.007 |  |  |  |  |  |
| **CY177** | 0.002 | 0.003 | 0.007 | 0.004 | 0.015 | 0.015 | 0.005 | 0.020 | 0.012 | 0.009 | -0.005 | 0.029 | 0.006 | 0.007 | 0.029 | 0.011 | 0.037 | 0.015 | -0.012 | 0.026 | 0.030 | 0.015 | 0.011 | 0.039 | 0.017 | 0.015 | 0.009 | 0.051 | 0.002 | 0.008 |  |  |  |  |
| **CY136** | 0.015 | 0.013 | 0.004 | 0.001 | 0.003 | 0.010 | 0.011 | 0.029 | 0.008 | 0.029 | -0.003 | 0.014 | 0.005 | 0.010 | 0.021 | 0.029 | 0.042 | 0.005 | -0.005 | 0.020 | 0.013 | 0.023 | 0.004 | 0.020 | 0.005 | 0.005 | 0.003 | 0.082 | 0.014 | 0.001 | 0.001 |  |  |  |
| **CY151** | 0.042 | 0.031 | 0.044 | 0.026 | 0.026 | 0.044 | 0.050 | 0.056 | 0.028 | 0.035 | 0.051 | 0.019 | 0.029 | 0.024 | 0.073 | 0.049 | 0.063 | 0.037 | 0.024 | 0.045 | 0.027 | 0.056 | 0.047 | 0.047 | 0.025 | 0.023 | 0.026 | 0.108 | 0.038 | 0.043 | 0.038 | 0.027 |  |  |
| **CY145** | 0.034 | 0.015 | 0.028 | 0.017 | 0.039 | 0.036 | 0.023 | 0.044 | 0.031 | 0.035 | 0.011 | 0.030 | 0.023 | 0.027 | 0.040 | 0.031 | 0.042 | 0.038 | 0.009 | 0.043 | 0.040 | 0.042 | 0.020 | 0.035 | 0.021 | 0.015 | 0.027 | 0.102 | 0.039 | 0.026 | 0.032 | 0.021 | 0.033 |  |
| **CY33** | 0.031 | 0.022 | 0.009 | 0.002 | 0.025 | 0.013 | 0.025 | 0.042 | 0.033 | 0.039 | 0.012 | 0.042 | 0.016 | 0.013 | 0.031 | 0.024 | 0.049 | 0.036 | 0.017 | 0.022 | 0.033 | 0.034 | 0.015 | 0.032 | 0.009 | 0.010 | 0.024 | 0.069 | 0.029 | 0.007 | 0.021 | 0.014 | 0.033 | 0.033 |

| **Buffalo** | **CY7** | **CY16** | **CY92** | **CY109** | **CY157** | **CY103** | **CY15** | **CY98** | **CY129** | **CY142** | **CY141** | **CY138** | **CY148** | **CY126** | **CY149** | **CY139** | **CY123** |
| --- | --- | --- | --- | --- | --- | --- | --- | --- | --- | --- | --- | --- | --- | --- | --- | --- | --- |
| **CY16** | 0.022 |  |  |  |  |  |  |  |  |  |  |  |  |  |  |  |  |
| **CY92** | 0.055 | 0.024 |  |  |  |  |  |  |  |  |  |  |  |  |  |  |  |
| **CY109** | 0.088 | 0.075 | -0.006 |  |  |  |  |  |  |  |  |  |  |  |  |  |  |
| **CY157** | 0.034 | 0.011 | -0.059 | -0.011 |  |  |  |  |  |  |  |  |  |  |  |  |  |
| **CY103** | 0.071 | 0.055 | -0.088 | 0.021 | -0.036 |  |  |  |  |  |  |  |  |  |  |  |  |
| **CY15** | 0.064 | 0.061 | 0.006 | 0.065 | -0.013 | 0.008 |  |  |  |  |  |  |  |  |  |  |  |
| **CY98** | 0.037 | 0.014 | -0.048 | 0.026 | -0.024 | -0.033 | -0.014 |  |  |  |  |  |  |  |  |  |  |
| **CY129** | 0.052 | 0.048 | -0.037 | 0.041 | -0.030 | -0.012 | 0.020 | -0.025 |  |  |  |  |  |  |  |  |  |
| **CY142** | 0.132 | 0.113 | -0.003 | 0.069 | 0.005 | 0.059 | 0.054 | 0.039 | 0.055 |  |  |  |  |  |  |  |  |
| **CY141** | 0.093 | 0.105 | 0.012 | 0.053 | -0.015 | 0.012 | 0.015 | 0.015 | 0.009 | 0.005 |  |  |  |  |  |  |  |
| **CY138** | 0.031 | 0.038 | -0.049 | 0.025 | -0.032 | -0.034 | -0.009 | -0.036 | -0.030 | 0.040 | 0.011 |  |  |  |  |  |  |
| **CY148** | 0.027 | 0.013 | -0.060 | 0.021 | -0.045 | -0.041 | -0.006 | -0.042 | -0.046 | 0.032 | -0.004 | -0.038 |  |  |  |  |  |
| **CY126** | 0.039 | 0.026 | -0.023 | 0.030 | -0.028 | -0.032 | -0.014 | -0.021 | -0.003 | 0.065 | 0.005 | -0.008 | -0.025 |  |  |  |  |
| **CY149** | 0.063 | 0.069 | -0.008 | 0.067 | -0.032 | 0.011 | 0.003 | 0.004 | -0.029 | 0.046 | -0.006 | -0.003 | -0.045 | -0.010 |  |  |  |
| **CY139** | 0.041 | 0.042 | 0.077 | 0.114 | 0.061 | 0.070 | 0.064 | 0.045 | 0.075 | 0.162 | 0.118 | 0.057 | 0.041 | 0.047 | 0.073 |  |  |
| **CY123** | 0.035 | 0.029 | 0.054 | 0.072 | 0.029 | 0.058 | 0.061 | 0.041 | 0.053 | 0.126 | 0.097 | 0.038 | 0.022 | 0.015 | 0.068 | 0.059 |  |
| **CY133** | 0.044 | 0.045 | 0.016 | 0.060 | 0.007 | 0.028 | 0.028 | 0.015 | 0.028 | 0.093 | 0.079 | 0.016 | 0.006 | 0.004 | 0.037 | 0.055 | 0.006 |
